# Towards a state-space geometry of neural responses to natural scenes: A steady-state approach

**DOI:** 10.1101/705376

**Authors:** Bruce C. Hansen, David J. Field, Michelle R. Greene, Cassady Olson, Vladimir Miskovic

## Abstract

Our understanding of information processing by the mammalian visual system has come through a variety of techniques ranging from psychophysics and fMRI to single unit recording and EEG. Each technique provides unique insights into the processing framework of the early visual system. Here, we focus on the nature of the information that is carried by steady state visual evoked potentials (SSVEPs). To study the information provided by SSVEPs, we presented human participants with a population of natural scenes and measured the relative SSVEP response. Rather than focus on particular features of this signal, we focused on the full state-space of possible responses and investigated how the evoked responses are mapped onto this space. Our results show that it is possible to map the relatively high-dimensional signal carried by SSVEPs onto a 2-dimensional space with little loss. We also show that a simple biologically plausible model can account for a high proportion of the explainable variance (∼73%) in that space. Finally, we describe a technique for measuring the mutual information that is available about images from SSVEPs. The techniques introduced here represent a new approach to understanding the nature of the information carried by SSVEPs. Crucially, this approach is general and can provide a means of comparing results across different neural recording methods. Altogether, our study sheds light on the encoding principles of early vision and provides a much needed reference point for understanding subsequent transformations of the early visual response space to deeper knowledge structures that link different visual environments.

## Introduction

On any given day, we receive a stream of visual information that is sampled from the environment in the form of retinal images. Exactly how the early visual system enables unique neural representations from this onslaught of visual information is a long-standing question in systems neuroscience. The last two decades have provided novel insights into the early visual encoding of real world stimulus images (“natural scenes”) in non-human vertebrates at levels of analysis ranging from single-units [1–9] to local population measures [10–12], and in humans using macro-scale measures such as EEG or fMRI [13–18] and psychophysics [19–24]. Together, such methods have contributed to a better understanding of how information is transformed along the visual pathway as well as why it is coded as it is.

Each of these techniques has both advantages and disadvantages and each reveals unique spatiotemporal features of the visual signal. EEG measures have the advantage that they are non-invasive, have good temporal resolution and can be relatively inexpensive. Despite their appeal, there remains considerable debate regarding the information that is provided by these signals. Studies with visual evoked potentials (VEPs) have focused on particular features of the response (e.g., N75, P100) and the effects that different stimulus conditions have on shaping the morphology of specific components. However, by focusing on particular features and not the signal as a whole, we feel that such an approach misses important properties of the signal. We believe that, rather than focusing on particular features, it is important to examine the full space of possible neural responses and consider how a population of responses falls within that space. By understanding the geometry of responses within this space, we believe that we can gain fundamental insights into the information available in these population measures.

In this study, we begin with a particular form of evoked cortical responses known as steady-state visual evoked potentials (SSVEPs), a well-established measure of visual responses in the early visual system (reviewed in [25–26]). Steady-state VEPs have a number of useful properties for measuring early visual responses in human observers [26]. Briefly, the SSVEP paradigm involves recording evoked potentials on the scalp while a participant views a stimulus that is modulated periodically at a particular frequency. If the stimulus drives a neural response that can be recorded on the scalp, the evoked potential will oscillate at the modulation frequency (and related harmonics) of the stimulus. An SSVEP can therefore be conceptualized by analogy to a steady-state response in a resonant circuit: by showing an observer an oscillating stimulus at a given frequency, the resulting electrical potentials entrain to the carrier frequency and remain stable in amplitude and phase [27–28]. We focus on SSVEPs because they allow us to collect neural response data with a signal to noise ratio (SNR) that is high enough to permit the recording of a relatively large number of images using relatively few stimulus repetitions.

The current study focuses on SSVEP responses to a broad population of natural scenes. A wide variety of studies have noted the importance of using ecologically relevant stimuli when probing sensory systems. The use of such stimuli allows us to observe the natural modes of activity of the visual system across the responses of different neural ensembles. In this study, we used a population of natural scenes as stimuli and determined the variety of brain responses that are produced by individual scenes as well as by repetitions of the same scene. Through the use of such stimuli, we will show that it is possible to use a relatively simple model of the early visual system to capture a high proportion of the explainable variance. By understanding how much of the neural response is driven by low level stimulus features, such models can allow one to deduce the amount of residual response variance that might be attributed to higher level factors.

The goal of the current study is to map and model the relative population responses that are generated by a set of natural scenes. Rather than focus on particular features of the neural response profile, we utilize a state-space approach. As we will show, one of the advantages of the SSVEP paradigm is that the output is low-dimensional, which allows us to consider a relatively simple state-space framework for understanding how images are organized by the early visual system. The state-space framework is a geometrical approach that considers the set of responses that a system produces in relation to the space of all possible responses. This geometric distribution of responses can then be understood in accordance with the distribution of images that have been projected to different encoding spaces (such as those defined by visual filter outputs). This general approach has been used in theories of sparse coding [29] and the non-linear behavior of visual neurons [30–31]. By focusing on the full state-space geometry of the responses produced by an evoked potential (rather than simple features of the response), our experiments will show that it is possible to provide both a rational model of the signal as well as to provide an estimate of the information carried by that signal.

We addressed the above across three experiments. In Experiment 1, we presented participants with a large set of natural scene images to measure the corresponding SSVEP-defined neural state-space. We assessed the reliability of that space by repeating images both within-and across experimental sessions, and quantified the reliability by estimating the mutual information between the SSVEP signal and each individual image. We replicated and extended these results with a larger set of images in Experiment 2, enabling us to better capture the boundaries of SSVEP state-space. Modeling revealed that 73% of the explainable variance in the SSVEP state-space could be accounted for by a Fourier filter-power model, a biologically plausible model of early visual processing. Experiment 3 then provided causal support for the modeling results. Overall, the techniques we describe here allow us to quantify the information provided by SSVEPs and to quantitatively characterize the organization of individual images by evoked potentials. The low-dimensional nature of the SSVEP state-space, while coarse, provides sufficient information for testing and contrasting theories of early visual processing. Moreover, the state-space framework is general, meaning that responses obtained from any neural recording technique (direct or indirect) can be projected into that space, thereby enabling comparative analyses of common stimulus sets across different types of recordings.

## Experiment 1

### Method

#### Apparatus

All stimuli were presented on a 23.6” VIEWPixx/EEG scanning LED-backlight LCD monitor with one ms black-to-white pixel response time. Maximum luminance output of the display was 100 cd/m^2^, with a frame rate of 120 Hz and resolution of 1920 × 1080 pixels. Single pixels subtended .0362° of visual angle as viewed from 36 cm. Head position was maintained with an Applied Science Laboratories (ASL) chin rest.

#### Participants

A total of 20 participants were recruited for this experiment. Of those, 3 failed to complete both recording sessions and 4 failed to produce signal to noise ratios (SNRs) at any electrode that exceeded chance SNR (measured on a participant-by-participant basis, described later). The age of the remaining 13 participants (4 female, 10 right-handed) ranged from 18-31 (median age = 20). All participants had normal (or corrected to normal) vision as determined by standard ETCRS acuity charts, gave Institutional Review Board-approved written informed consent before participating, and were compensated for their time.

#### Stimuli

Stimuli were selected from a large database of real-world scenes consisting of 2500 photographs that varied in content from purely natural to purely carpentered (both indoor and outdoor), with various mixtures of natural/carpentered environments in between. The images were largely sampled from several existing databases [20, 32–33], with several hundred sampled from Google Images (copyright-free). Selection criteria included 1) images that were in focus at all depths, 2) had a minimum pixel dimensions between 512 and 1024, and 3) were largely devoid of people or faces. All images were then cropped to 512 × 512 pixels and converted to grayscale using the standard weighted sum conversion in Matlab.

Stimuli were selected by randomly sampling 150 images from the image database. All stimuli subtended 18.5° of visual angle, and were made to possess the same root mean square (RMS) contrast (0.2) and mean pixel value (127) as described in **Appendix 1**. All images were fit with a circular linear edge-ramped window (512-pixel diameter, ramped to the pixel mean) to obscure the square frame of the images, thereby ensuring contrast changes at the boundaries of the image were not biased to any particular orientation [20, 34].

#### Procedure

The experiment consisted of two recording sessions, each lasting ∼55 minutes. Within each session, all 150 stimuli were presented once in each of two sequential blocks, with a random order within each block, resulting in a total of four repetitions per image over both recording sessions. Each trial began with a 3000 ms blank (mean gray) screen, followed by a 6000 ms stimulus interval. During that interval, the stimulus image was contrast-modulated at a rate of 5 Hz with a *sinusoidal* temporal profile^1^. Participants were engaged with a distractor task at fixation, which consisted of detecting a luminance change (black to white) of a 4 × 4-pixel square placed at the center of the stimulus image. The luminance change occurred on 50 random trials, and the images for those trials were not considered in subsequent analyses (an additional 50 images were used on those trials). Thus, a total of 350 images (300 experimental, and 50 luminance change) images were presented per session. Participants reported when a luminance change occurred via gamepad response, and were given the opportunity to rest every 50 trials.

#### EEG Recording and Processing

Continuous EEGs were recorded in a Faraday chamber using Electrical Geodesics Incorporated’s (EGI; Philips Neuro) Geodesic EEG acquisition system (GES 400). All EEGs were obtained by means of Geodesic Hydrocel sensor nets consisting of a dense array of 128 channels (electrolytic sponges). The on-line reference was at the vertex (Cz), and the impedances were maintained below 50 kΩ (EGI amplifiers are high-impedance amplifiers). All EEG signals were amplified and sampled at 1000 Hz. The digitized EEG waveforms were band-pass filtered offline from 0.1 Hz to 50 Hz to remove the DC offset and eliminate 60 Hz line noise. Finally, each trial event was tagged via photodiode response to a small white square that flashed in the upper left-hand corner of the display (obscured from participant’s view) at the start of each trial. Tagging trials in this manner eliminated the offset time and clock drift between the acquisition computer and the experiment station computer.

To remove onset transients from the analysis, the first 1000 ms of the stimulus interval was removed. Thus, all continuous EEGs were segmented into 5000 ms waveforms corresponding to the last 5000 ms of the stimulus interval. Segments that contained eye-movements, eye-blinks, or transients greater than ± 250 µV (fewer than 7% of the trials on average) were flagged but were found to have no impact on the stimulus fundamental frequency (or its harmonics) and were therefore included in all subsequent analyses. Topographic plots were generated for all experimental conditions using EEGLAB (Delorme & Makeig, 2004) version 13.5.4b in Matlab (ver. R2017a).

#### Electrode Selection

Electrode selection was carried out in a data-driven manner via significance testing. First, we calculated the SNR for each trial epoch by fast Fourier transforming each electrode’s EEG waveform, and then divided the amplitude at the fundamental frequency (5Hz) by the average of the neighboring frequencies (4.4 to 4.8 Hz and 5.2 to 5.6 Hz), referred to here as the *noise denominator*. In order to compare the observed SNR to what could be expected by chance, we estimated the null SNR for each electrode across all epochs by randomly sampling two noise denominators and taking their ratio. This process was repeated 5000 times to generate a null SNR distribution for each electrode. To determine which electrodes produced SNRs that were significantly different (p < 0.05) from their corresponding null SNR distribution, right-tail z-tests were run for all SNRs for each electrode (across all trials) against that electrode’s null SNR distribution. The electrode with the largest effect size (“best electrode”) was automatically included in subsequent analyses. To test for other electrodes with averaged SNRs on par with the “best electrode”, we tested the best electrode’s SNR distribution against all other significant electrode SNR distributions (left-tail z-tests), and those that were not significantly different from the best electrode were also included in subsequent analyses. This method identified 3-8 electrodes across participants, and these were always located over the occipital pole. Participants were excluded from subsequent analyses if none of their electrode SNRs (averaged across trials) were different from their corresponding null SNR distributions, or if their SNRs were all below 2, as SNRs below that value were observed to result from a lack of entrainment of the 5Hz fundamental to the phase of the sinusoidal stimulus modulation.

Lastly, the final time series data were generated by averaging epochs across all selected electrodes to yield a single 5000 (time points) × 300 (trials) matrix for each participant and for each recording session.

### Results

#### SSVEP Signal Characteristics

To measure signal quality for each image, all repetitions for each image were averaged together in the time domain, thereby reducing the two 5000 × 300 matrices to a single 5000 × 150 matrix which was used to calculate an SNR spectrum for each trial (**Figure 1b**). Each SNR spectrum was calculated by measuring the SNR for each frequency in the Fourier amplitude spectrum of each SSVEP. The size of the windows used to define the noise denominator was the same as that described in the Electrode Selection section. Each trial’s SNR spectrum was then sampled at whole integer frequencies and averaged across trials. This process was repeated for each participant, and then averaged across participants and plotted in **Figure 1b**. The experimental paradigm yielded strong signal strength at the fundamental frequency (5 Hz) as expected, as well as the next three harmonics that reflect nonlinear responses that were also entrained to the stimulus modulation. For a more complete view of each entrained signal, the averaged time series data were filtered (in turn) at the fundamental and each of the three harmonics to extract each frequency’s phase which was then plotted in polar form along with each frequency’s SNR (**Figure 1c**). The results show that three of the four frequencies have the majority of their phase angles falling within ±30**°** of a central angle, with the 10Hz harmonic showing a similar phase angle tuning centered on mean angles that differ by ∼180**°. Figure 1c** therefore illustrates that the periodic modulation was successful in entraining the fundamental and harmonic frequencies because an un-entrained oscillation would yield a uniform distribution of phase angles.

**Figure 1.**
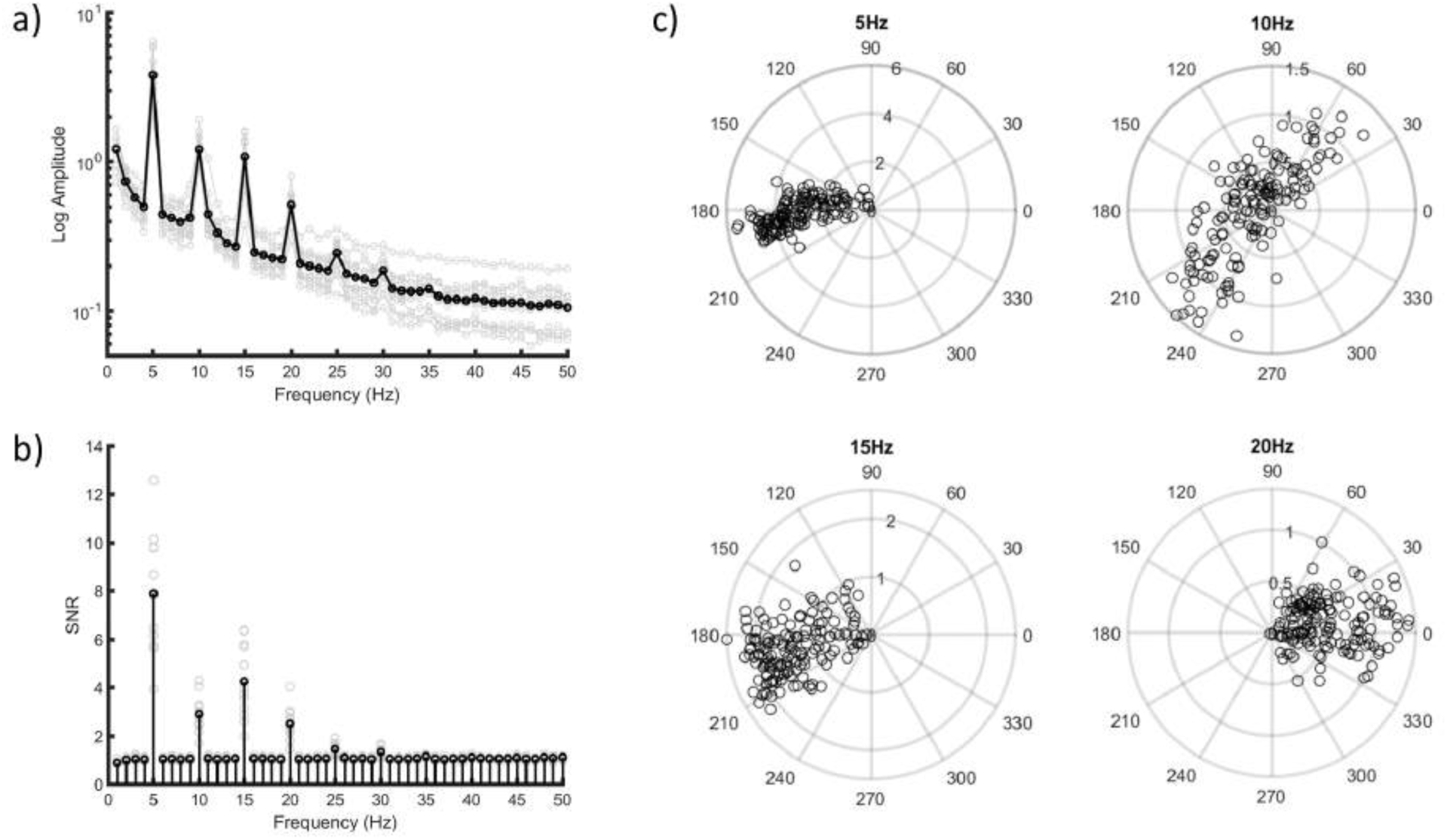
Experiment 1 signal characteristics. (a) Amplitude spectrum resulting from averaging across images and participants (black trace). Each light gray trace shows each participant’s amplitude spectrum averaged across images. (b) Signal-to-noise (SNR) spectrum calculated from the across participant and image average shown in black, with gray symbols showing each participant’s SNR averaged across images. (c) Polar plots showing participant and image averaged SNR (radial axis) and phase angle (theta axis) for the fundamental frequency (5Hz) and the three most prominent harmonic frequencies.

#### Principal Component Analysis

The results illustrated in **Figure 1** show that there are a relatively small number of Fourier components in the SSVEP response. If we think of the first four Fourier components, each with real and imaginary components (or amplitude and phase), this suggests that the denoised signal is no more than 8-dimensional. However, the dimensionality may be much lower if some of those dimensions are highly correlated. To measure the true dimensionality of SSVEP state-space, we turn to principal component analysis (PCA).

Having only four stimulus modulated frequencies in the SSVEP signal (e.g., **Figure 1a-b**) means that the rest of the energy in the signal was not modulated by the stimulus and can therefore be treated as noise. We filtered the time series data to contain the fundamental plus the three harmonic frequencies. That particular compound-frequency signal implies that the dimensionality is no more than eight (four frequencies each with a sine and cosine component), but could be significantly less. Further, because the amplitude of each frequency is the sum of signal and noise, we normalized each frequency’s waveform peak to its corresponding SNR on a trial-by-trial basis. We then submitted the 5000 × 150 filtered and SNR-normalized data matrix to PCA. The first three principal components (PCs) were found to account for 97% of the variance in the data, with the first PC accounting for most (92%, see Figure 2a). To evaluate whether the four stimulus induced frequencies contributed to each of the PC basis functions (i.e., the PC scores), we submitted each PC’s basis function to Fourier analysis. The results of that analysis revealed that all four frequencies contribute to each PC dimension (**Figure 2b-d**). To assess the reliability of the first three PC dimensions, PCA was conducted on the SSVEP matrices for each repetition (on a participant-by-participant basis). Next, each PC’s eigenvector was correlated with its corresponding eigenvector across each repetition. We found that only PC1 and PC2 produced statistically significant correlations across stimulus repetition (across participants, the correlation cofficients for PC1 ranged between 0.45 – 0.72, 0.18 – 0.35 for PC2, and −0.08 to 0.06 for PC3). Thus, only PCs 1 and 2 captured variance in a reliable manner across stimulus repetition, and were included in all subsequent analyses. To visualize SSVEP state-space, we plotted the joint distribution of images defined by the first two eigenvectors (i.e., the rotation of each image’s SSVEP to each PC’s basis function) (**Figure 2e-f**). The lack of symmetry in this space implies that these first two eigenvectors are not independent. Because each component is a linear combination of the four frequencies, the data points have been color coded according to the average SNR-normalized amplitude **(Figure 2e)** and averaged phase angle **(Figure 2f)**. In order to express phase angle linearly, we used circular averaging [35]. As a result, we see that the first PC is extracting signal magnitude (R^2^ = 0.95, p < 0.001), with the second partially coding for phase (circular-linear R^2^ = 0.48, p < 0.001).

**Figure 2.**
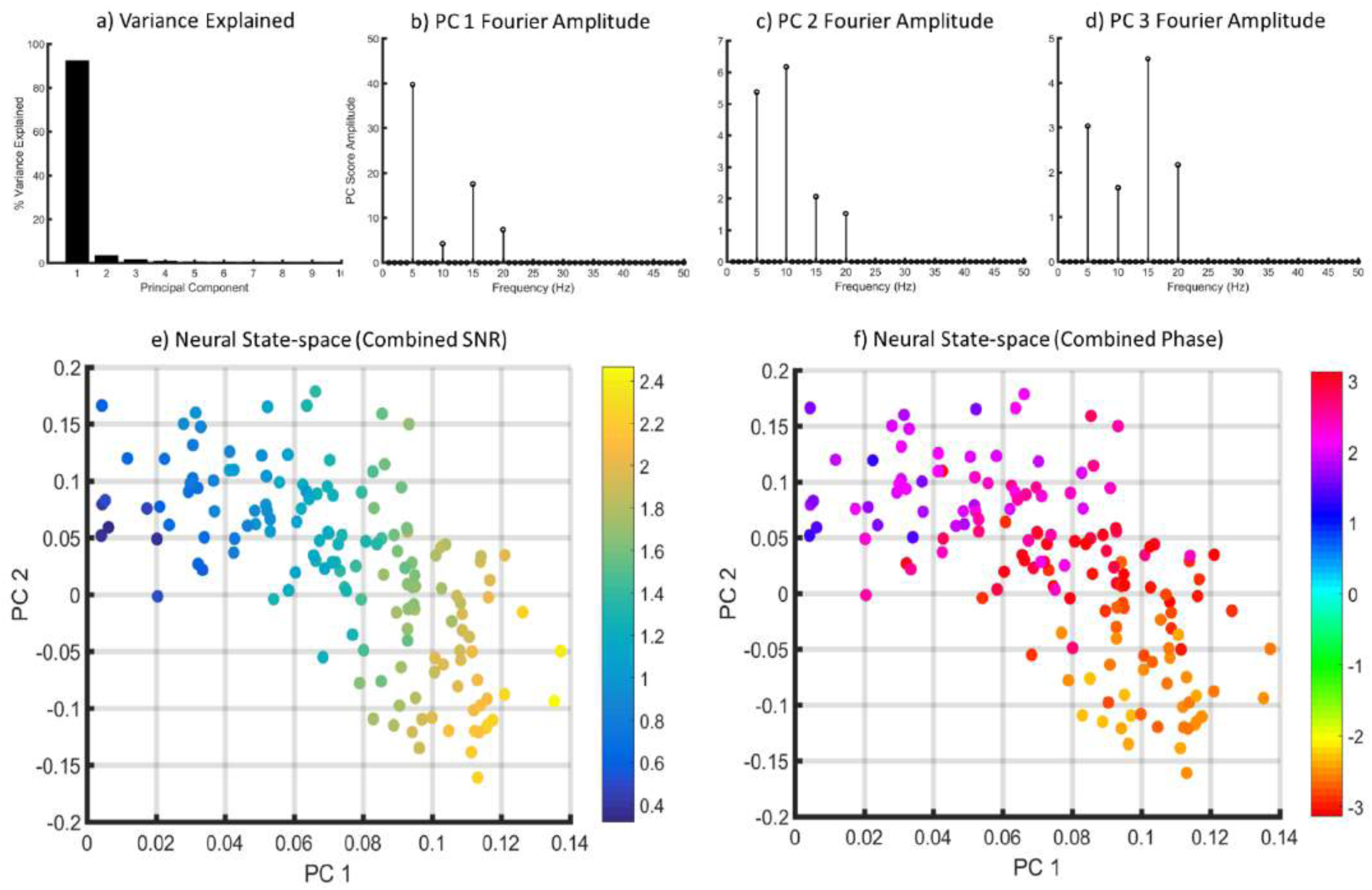
Experiment 1 Principal Component (PC) Analysis results. (a) Percentage of variance explained by the first 10 PCs. (b-d) Amplitude spectrum for each of the first three PC basis functions (i.e., PC scores) showing that each PC contains amplitude at the fundamental and next three harmonic frequencies. Note that the scales of those plots are different. (e) Principal component coefficients 1 and 2 for the 150 images. Here each data point is an image in neural state-space and has been color coded according to the average SNR of the fundamental and next three harmonic frequencies. (f) The exact same space color coded according to the circular average of the phase angles of the fundamental and harmonic frequencies.

Next, we examined how the images are organized along the eigenvector axes of the SSVEP state-space. **Figure 3** (bottom) shows images ordered according to PC1’s eigenvector coefficients from lowest (left) to highest (right) (rows are in arbitrary order). The ordering of images according to the coefficients for PC2 are shown in the Supplementary Materials section (Figure S1). We also generated topographic plots of the averaged SNR for each electrode for each of the five bins (**Figure 3**, top). As noted in the method section, SNRs were greatest at electrodes over the occipital pole and here we see that SNR increases in magnitude in proportion to PC1’s eigenvector coefficients at those electrodes. Interestingly, the organization of the images along PC1 bears a striking resemblance to the spatial principal components of scenes as reported by [36] and seem to be organized in terms of increasing contrast energy at high spatial frequencies. This observation was verified by measuring the amount of Fourier power of the stimuli with a wide range of log-Gabor filters (detailed in **Appendix 2**) tuned to different spatial frequencies (SFs) (0.05, 0.125, 0.25, 0.50, 1.0, 2.0, 4.0, 6.0, and 8.0 cycles per degree, cpd) and orientations (0°-165° in steps of 15°) – **Figure 4**. The resulting filter spectra show a clear transition from being largely dominated by lower SFs at smaller PC1 eignevector coefficients to largely high SF dominated at larger eigenvector coefficients. We will return to this observation in the Information Analysis and Modeling sections of this study.

**Figure 3.**
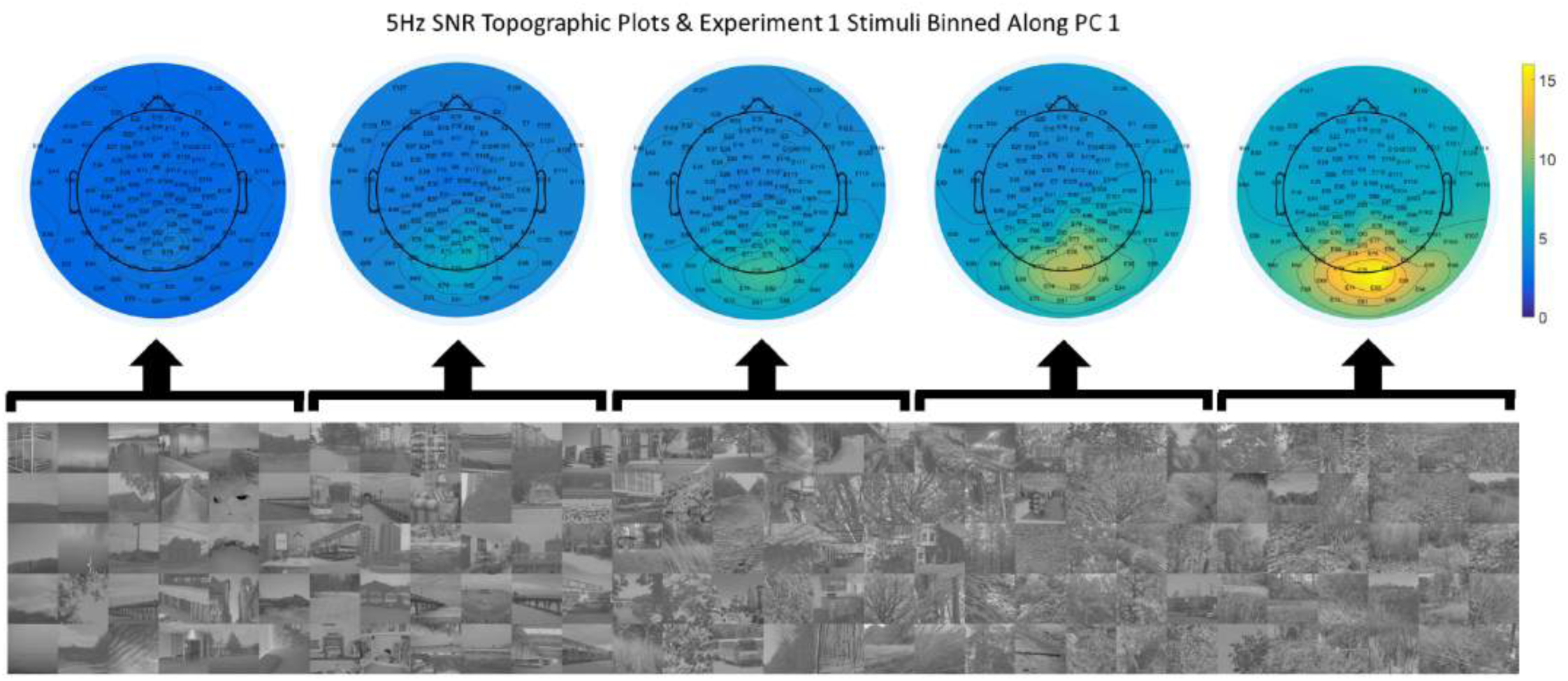
Experiment 1. Top: topographic plots of SNR (averaged across participants) for the 5 Hz fundamental for sets of images organized along PC1 (small coefficients on the left and large coefficients on the right). Bottom: Stimuli binned along PC1 (x-axis of the image array). To facilitate a visual presentation, the images were sorted according to the participant-averaged PC1 coefficients and binned into 5 sets with 30 images per bin (thus there is no inherent meaning to the y-axis of the image array).

**Figure 4.**
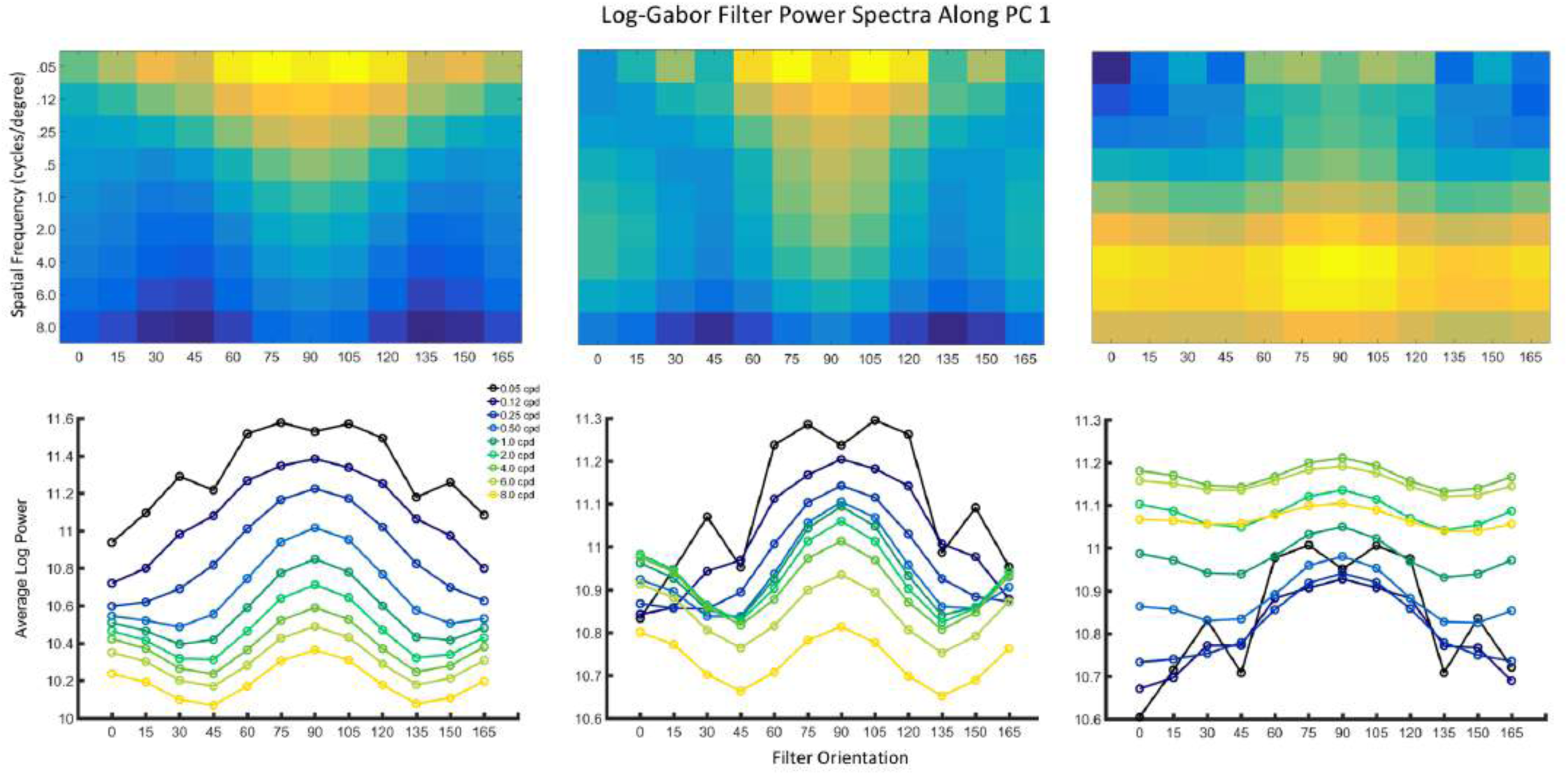
Fourier filter-power analysis of the stimuli used in Experiment 1, organized along PC1 from left-to-right (smaller coefficients to large coefficients). The top row shows the Fourier power for each of the 108 filters. Rows show the spatial frequency (SF) in cycles per degree (cpd) peak of each filter and the columns show the peak orientation (in degrees) of each filter. The bottom row replots the filter power spectra shown above in a standard line plot format.

## Experiment 2

The results of Experiment 1 show that the SSVEP signal can be described in a low-dimensional space whose dimensions systematically code for visual features known to be extracted in early visual processing. However, in order to measure that space with reasonable stability, each image was repeated four times, thereby placing an upper limit on the number of unique images included in that experiment. As a result, our sampling of the natural scene defined SSVEP state-space was relatively sparse, possibly blurring the boundaries of the space. Here, we sought to characterize the state-space boundaries observed in Experiment 1 by probing that space with more images (700 in total). To keep the experiment within a reasonable time limit, each image was presented only once. However, we aimed to ‘regain’ signal quality at the image level by averaging the SSVEP time series data across a reasonably large number of participants.

### Apparatus

Same as in Experiment 1.

### Participants

A total of 25 participants were recruited for this experiment. Of those, 2 failed to complete both recording sessions and 5 failed to produce SNRs at any electrode that exceeded chance SNR (detailed in Experiment 1). The age of the remaining 18 participants (8 female, 16 right-handed) ranged from 17 to 23 (median age = 18). All participants had normal (or corrected to normal) vision as determined by standard ETCRS acuity charts. All participants gave Institutional Review Board-approved written informed consent before participating and were compensated for their time.

### Stimuli

The stimuli consisted of the same 150 images from Experiment 1 plus 550 additional images randomly sampled from our 2500 image database. All stimuli were prepared as described in Experiment 1.

### Procedure

The stimuli were randomly assigned to one of two recording sessions and were randomly interleaved within each recording session (350 images per session, with each session lasting ∼55 min). The trial sequence and contrast modulation were identical to Experiment 1. Participants were engaged with a distractor task at fixation, which consisted of detecting a color change (blue to red or green) of a 4 × 4 pixel square placed at the center of the stimulus image. When the color changed during each trial, and whether or not a color change occurred at all was determined randomly. Participants reported the color changes via gamepad response and were given the opportunity to rest every 50 trials.

The EEG recording details, data processing pipeline, and electrode selection routine were identical to Experiment 1. As in Experiment 1, artifact trials (no more than 8% across all participants) were found to have no influence on the fundamental frequency and were therefore included in all subsequent analyses. Thus, each participant’s data consisted of an electrode-averaged time series matrix that was 5000 (time points) × 700 (stimuli). As in Experiment 1, the number of included electrodes ranged from 3-8 across participants. All participant data matrices were then averaged and submitted for analysis.

### Results

#### SSVEP Signal Characteristics

We calculated the SNR spectrum as in Experiment 1, except here we used the participant-averaged 5000 × 700 data matrix. This approach yielded strong signal strength at the fundamental frequency (5Hz), as well as the next three harmonics, consistent with Experiment 1 (**Figure 5b**). Next, the averaged time series data were filtered (in turn) at the fundamental and each of the harmonics to extract each frequency’s phase angle which was then plotted in polar form along with each frequency’s SNR (**Figure 5c**). The results are consistent with Experiment 1 in that the phase angles largely fall between ±30**°** of a central angle (excepting the 20 Hz harmonic, which mostly fell between ±60**°** of its central angle).

**Figure 5.**
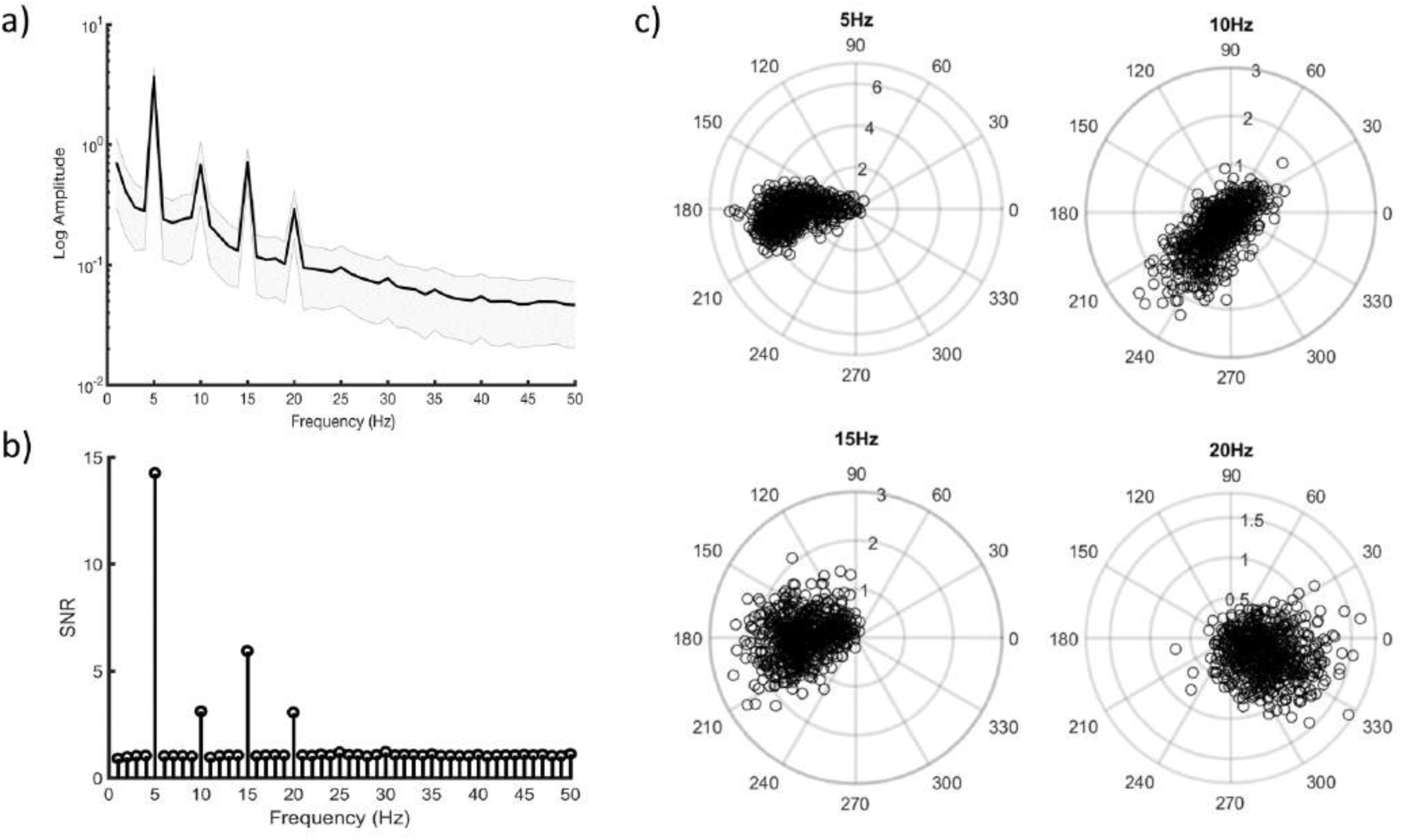
Experiment 2 signal characteristics. (a) The amplitude spectrum of the participant averaged time series data (black trace) bounded by 95% confidence intervals (shaded gray) across all 700 stimulus trials. (b) The SNR spectrum calculated from the participant averaged time series data. (c) Polar plots showing participant and image averaged SNR (radial axis) and phase angle (theta axis) for the fundamental frequency (5Hz) and the three most prominent harmonic frequencies.

#### Principal Component Analysis

The participant-averaged time series matrix was filtered to contain the fundamental plus the three harmonic frequencies, and SNR-normalized as in Experiment 1. We then submitted the resulting 5000 × 700 data matrix to PCA. The first three PCs were found to account for 96% of the variance in the data, with the first PC accounting for most (90%), similar to Experiment 1. Fourier analysis of the first three PC basis functions revealed patterns that were virtually identical to those shown in **Figure 2b-d**. The coefficients from the first two eigenvectors were plotted against one another (**Figure 6**) and show a non-symmetric distribution as was observed in Experiment 1. **Figure 6** is color coded by average SNR-normalized amplitude and phase angle as they were in Experiment 1 (**Figure 2**) and shows the same SNR amplitude and phase angle relationship, specifically, the first PC is extracting signal amplitude (R^2^ = 0.96, p < 0.001), with the second partially coding for phase (circular-linear R^2^ = 0.49, p < .001).

**Figure 6.**
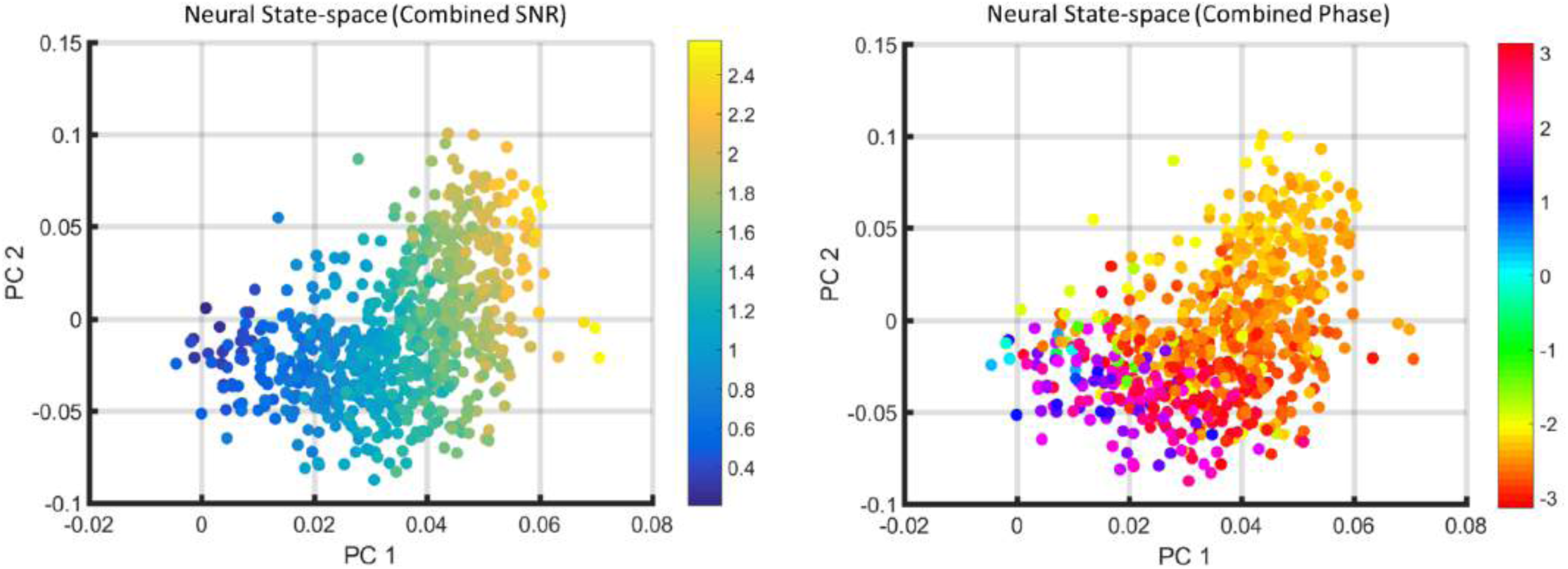
Left: Principal component coefficients 1 and 2 for the 700 images used in Experiment 2. Each data point is an image in neural state-space and has been color coded according to the average SNR of the fundamental and next three harmonic frequencies. Right: The exact same space color coded according to the circular average of the phase angles of the fundamental and harmonic frequencies.

A visual demonstration of how the images are organized along the eigenvector axes of the SSVEP neural response-space is provided in **Figure 7** (bottom). The ordering of images according to the coefficients for PC2 are shown in the Supplementary Materials section (Figure S2). We also generated topographic plots of the averaged SNR for each electrode (**Figure 7**, top). The organization of the images according to the eigenvector coefficients for PC1 is very similar to that observed in Experiment 1 (**Figure 3**), which was confirmed with regression between the Euclidean distances between PC1 coefficients of the corresponding images across both experiments (R^2^ = 0.65, p < .001).

**Figure 7.**
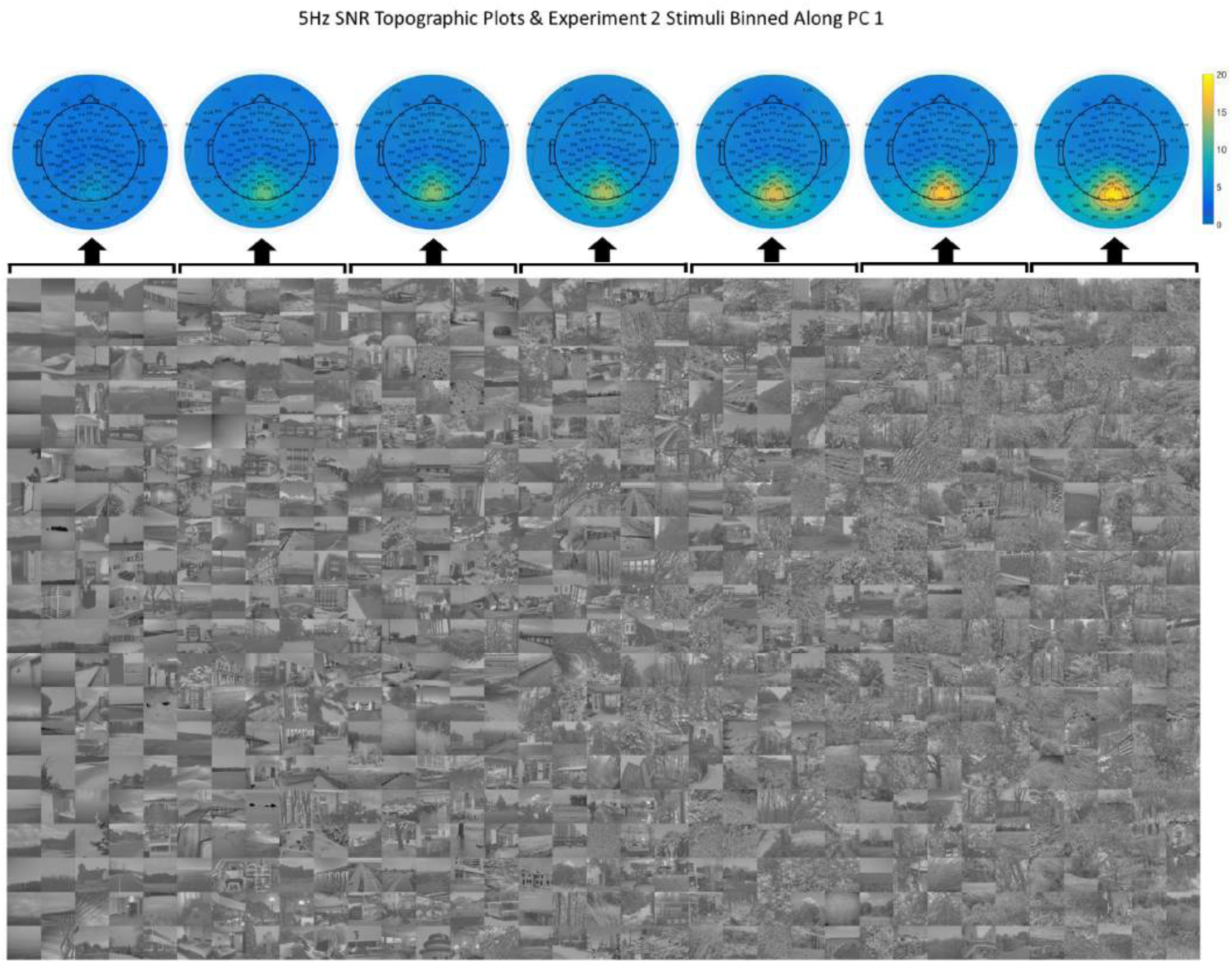
Experiment 2. Top: topographic plots of SNR (averaged across participants) for the 5 Hz fundamental for sets of images organized along PC1 (small coefficients on the left and large coefficients on the right) – each topographic plot corresponds to the 20 × 5 image bin directly below. Bottom: Stimuli binned along PC1. To facilitate a visual presentation, the images were sorted according to the participant-averaged PC1 coefficients (low to high) and binned into 7 sets with 100 images per bin (as with **Figure 3**, row membership is arbitrary).

### Information Analysis

The data from Experiments 1 and 2 allow us to provide an estimate of the information that is carried by the SSVEP signal with respect to our population of natural scenes. The information is a function of the reliability of the SSVEP response across repeated image presentations as well as the uniqueness of the response to each image. The analysis described below calculates the mutual information between our image set and the SSVEP responses from Experiment 1. It is important to first recognize the particular constraints imposed on these conclusions. First, we are describing the average information in the signal with respect to a particular population of natural scenes. These scenes are normalized to have the same RMS contrast and they extend 18.5 degrees centered on the fovea. Second, we are not attempting to calculate the true entropy of natural images (which would be much too difficult -i.e., see [37]). Rather, we are considering our set of images as a finite set of stimuli (150 images = 7.23 bits) and the information we calculate provides an estimate of how accurately a particular image can be identified given the SSVEP signal (the maximum possible is 7.23 bits). Third, we are also using the “best electrode” approach as described in the previous section and not using the information contained in the spatial distribution of activity across the different electrodes. Fourth, we use just four presentations of the stimulus to make an estimate of the variance of the response. Finally, we are making this estimate with only the first two principal components because the third PC accounted for only 1.76% of the variance and did not show clear relationship with repeated measures.

Given these limitations, there are a number of values that we can calculate. Each value provides only a rough estimate, but they provide broad insights into the reliability of the signal. The mutual information between two signals (X and Y) is defined as the difference between two entropies:

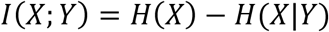

For this study, we define H(X) as the entropy of the SSVEP signal across the entire set of images. Intuitively, this quantity should reflect the spread of image points across a space defined by the first two PCs of SSVEP activity. To operationalize this, for each of the four presentations of each image, we computed the mean and covariance of the coefficients from the first two eigenvectors, and modeled each image as a multivariate elliptical Gaussian distribution. We then summed these Gaussians together, and divided by the total to create a probability distribution over PC space, *p* in a grid of 100 by 100, defined by the range of each principal component. Entropy was then computed as standard:

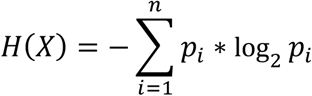

Where n is the number of cells in the grid^2^.

H(X|Y) is then defined as the entropy of the SSVEP signal conditioned on a particular image. For this study, we operationalized this as the probability density defined by the multivariate elliptical Gaussian fit to the four presentations of each single image. The mutual information value for each participant is the average mutual information over the 150 images.

As shown in **Figure 8a**, when there is a good deal of similarity in the SSVEP across the four image presentations of a given image, there is a higher resulting mutual information value. This is because the variability in PC locations across presentations is small, resulting in a smaller H(X|Y) relative to H(X). On the other hand, when SSVEP responses are variable across image presentations, higher variability in PC locations result in larger H(X|Y). In other words, the probability distribution for that image becomes more similar in size to that of the entire set of images, captured by H(X). This results in low mutual information between the SSVEP and the given image.

**Figure 8.**
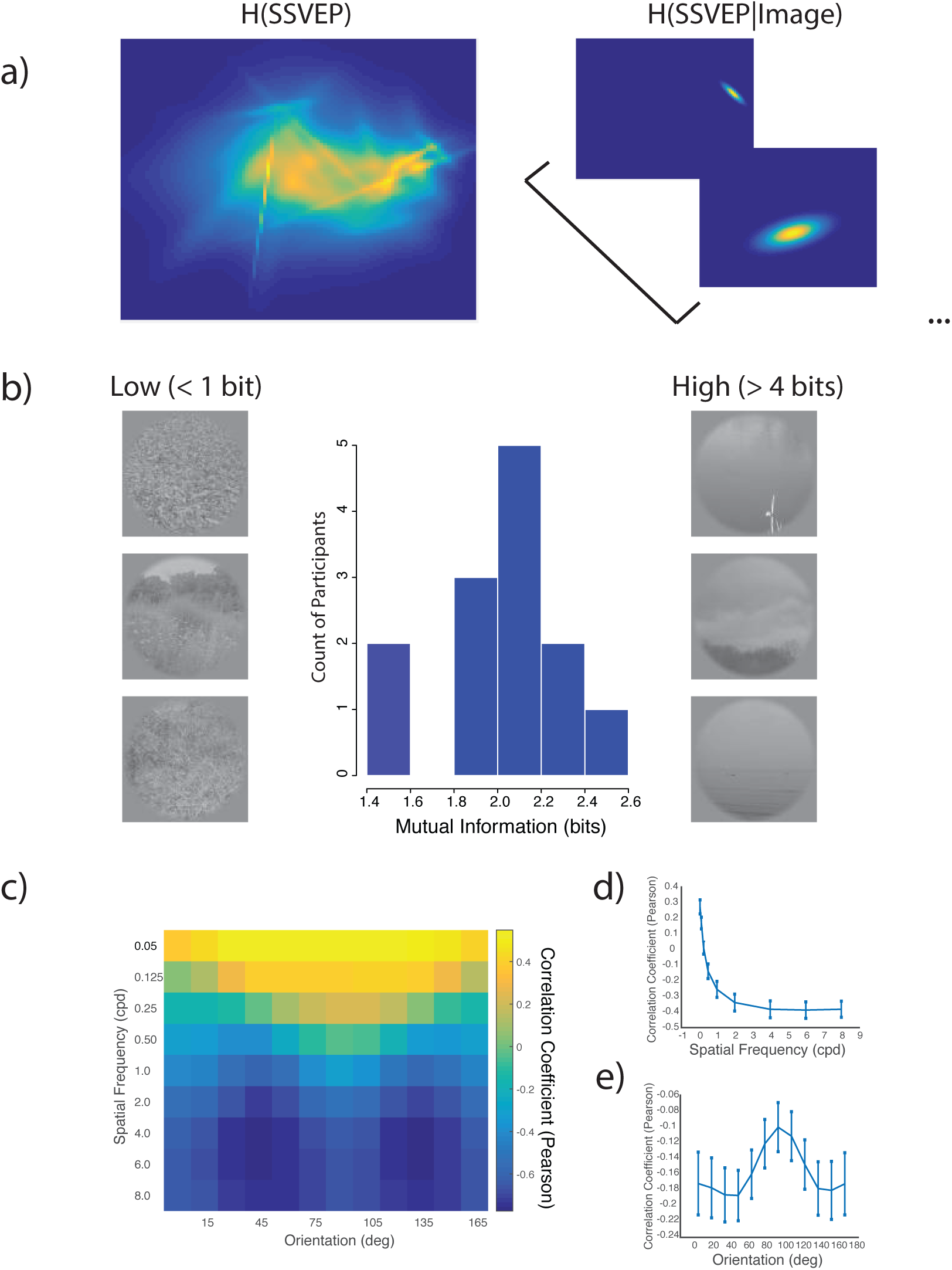
Mutual information between image and SSVEP signal. (a) Left: We fit multivariate Gaussian distributions to the four presentations of each of the 150 images in a space defined by the first two PCs. The entropy of the SSVEP signal (H(SSVEP)) was estimated by summing these 150 Gaussians. Right: The individual Gaussians constitute the image-defined entropy (H(SSVEP|Image)). The mutual information is the difference between these quantities. (b) Histogram of mutual information values across participant (mean=2.1 bits). Side images indicate images with particularly low-or high-mutual information with SSVEP signal. (c) Correlation of mutual information and Gabor filter parameters (spatial frequency and orientation). (d) Marginal correlation of spatial frequency and mutual information. (e) Marginal correlation of orientation and mutual information. For both (d-e), error bars indicate +/-95% confidence interval.

We found that the mutual information between the SSVEP responses and our image set is ∼2.1 bits (range across participants: 1.5 to 2.5 bits). This value gives us an estimate of how much information the SSVEP signal gives us about the specific image being presented. Additionally, we collapsed across participants and examined the distribution of mutual information across images. As shown in **Figure 8b**, we found that images with low mutual information with the SSVEP signal tended to have more high spatial frequency content than those with higher mutual information. We tested the generality of this observation by correlating the average mutual information value of each image with the filter power of the images at nine spatial frequencies and 12 orientations (detailed in the Results section of Experiment 1) (**Figure 8c**). We found that across all images, low spatial frequencies were associated with higher mutual information (R^2^ = 0.55, p < 0.05, collapsed across orientation, **Figure 8d**). By contrast, orientation (when collapsed across spatial frequency) was not linearly associated with mutual information (R^2^ = 0.03, n.s.). However, as shown in **Figure 8e**, this is because orientations around 90° were associated with higher mutual information than the oblique angles of 45° and 135°.

Again, we must emphasize that this measure of information is only a general estimate limited to the conditions we have used in this study, and most likely a lower bound. With only four presentations of each image, we are likely overestimating the variability of each image. A wider range of natural scenes (e.g., where the contrast is not constrained or the images are colored or larger) may likely provide a larger value. It is also likely that if we opened the data set to a much wider variety of scenes (e.g., gratings, abstract textures, etc.) that we would find that the SSVEP signal carries more information about the signal.

While it is worth knowing that SSVEP signals carry significant information regarding image content, this approach provides less insight about what kind information is carried in that signal. To explore this question, we developed a model of how the SSVEP signal is represented by the visual system, allowing us to determine what aspects of the stimulus predict the SSVEP responses. In the next section, we describe this model and demonstrate that it can predict a relatively high proportion of the response variance.

### Explaining the SSVEP Response Space

The analyses reported thus far point to spatial frequency being an important organizing factor along PC1 (e.g. **Figure 4**), as well as a modulator of mutual information between the stimulus and response (e.g. **Figure 8D**). This suggests that the mapping between image state-space and SSVEP response space may rely on a Fourier-power based encoding scheme, a well-justified model of the early visual system (V1 in particular) [38]. Here, we test this empirically by measuring the relationship between Euclidean distances between the filter power spectra of our stimuli and their corresponding distances in SSVEP response state-space.

Steady-state visual evoked potentials represent a global measure of the underlying neural operations at the circuit level, meaning that the entrained signal measured on the scalp likely stems from a summation of the underlying responses tuned to different image attributes. If the majority of the summation arises from early visual cortical processes (reviewed in [38]), then we can expect a good portion of the sum to be explained by contrast in different bands of spatial frequency and orientation. Therefore, to model the cortical response, we represented each image in terms of an array of filters selective to different positions, spatial frequencies and orientations inline with the sort of tuning that is found in area V1. Although it is certainly possible that the SSVEP signal carries higher level information, we believe a strong, initial approach is to see what can be explained by these low level features.

All stimuli in Experiments 1 and 2 were filtered using log-Gabor filters (detailed in **Appendix 2**) centered on nine different spatial frequencies and 12 orientations. The filters were set to have an SF bandwidth of 1.4 octaves (full width at half height) and an orientation bandwidth of 36° (full width at half height) [39–40]. Each image’s power spectrum was multiplied by each filter and summed and we then calculated the log of this sum. After this transformation, each image is represented as an array of 108 filter responses (log Fourier power), which can be used to construct an item-by-item Euclidean distance matrix, which can then be directly compared to the item-by-item Euclidean distances in the eigenvector-defined SSVEP state-space [41]. This process resulted in a 150×150 distance matrix for Experiment 1 and a 700×700 distance matrix for Experiment 2. We then calculated Euclidean distance matrices between each image’s location along each axis of the SSVEP state-space for Experiments 1 and 2 separately. To ensure that the distances reflected the difference in variance explained by PC 1 and 2, the eigenvector coefficients were first weighted by the square root of each PC’s corresponding eigenvalue. Regression analyses between filter output distances and state-space distances from each experiment resulted in only modest relationships (Expt 1: R^2^ = 0.32, p < .001; Expt 2: R^2^ = 0.30, p < .001). However, Experiments 1 and 2 both revealed that the first PC accounted for > 90% (in both experiments) of the variance in the SSVEP response space. While the eigenvectors were weighted by the square root of their corresponding eigenvalues in the above analysis, it is likely that the inclusion of PC2 added noise to the aforementioned analysis due to the small amount of variance that it explained. Further, the filter-power distances were driven by the log power of 108 filters, many of which may not have been instrumental in driving SSVEP entrainment.

To provide an analysis more suitable to the PC-defined SSVEP state-space, the log filter-power across all stimuli for each peak SF and orientation was regressed against PCs 1 and 2 separately for each experiment (i.e., we attempted to model each eigenvector dimension of the SSVEP state-space). The results using the PCs from Experiment 1 are plotted in **Figure 9a-b**. Interestingly, for both PCs, we see that most of the variance in the PC eigenvector coefficients is accounted for by filter-power at the higher spatial frequencies, with a bias at the oblique orientations. We then repeated this analysis for each participant. To build a full model that minimized multicollinearity across filters, we averaged together the power at the cardinal orientations (0° and 90°) and oblique orientations (45° and 135°) for each image (resulting in 18 predictors, 9 SFs each for cardinal and oblique orientations). The filter-power predictors were then entered into a multiple regression against PC1 eigenvector coefficients for each participant (based on the average of all four repetitions). We estimated upper and lower bounds of explainable variance given the noise inherent in the data according to **Appendix 3** on a participant-by-participant basis (**Figure 9c**). We then estimated the success of the model by taking the ratio of the model R^2^ and the upper bound of explainable variance. The model was able to account for 72.8% (SE = 2.17%) of the explainable variance in PC1 coefficients (**Figure 9c**), and 60% (SE = 4.5%) of explainable variance in PC2 coefficients (not shown). For both PCs, the higher SFs (≥4 cpd) accounted for an averaged ∼86% of the full model’s R^2^. Thus, a simple Fourier filter-power model can explain an impressive portion of the organization of SSVEP neural response-space along both PCs.

**Figure 9.**
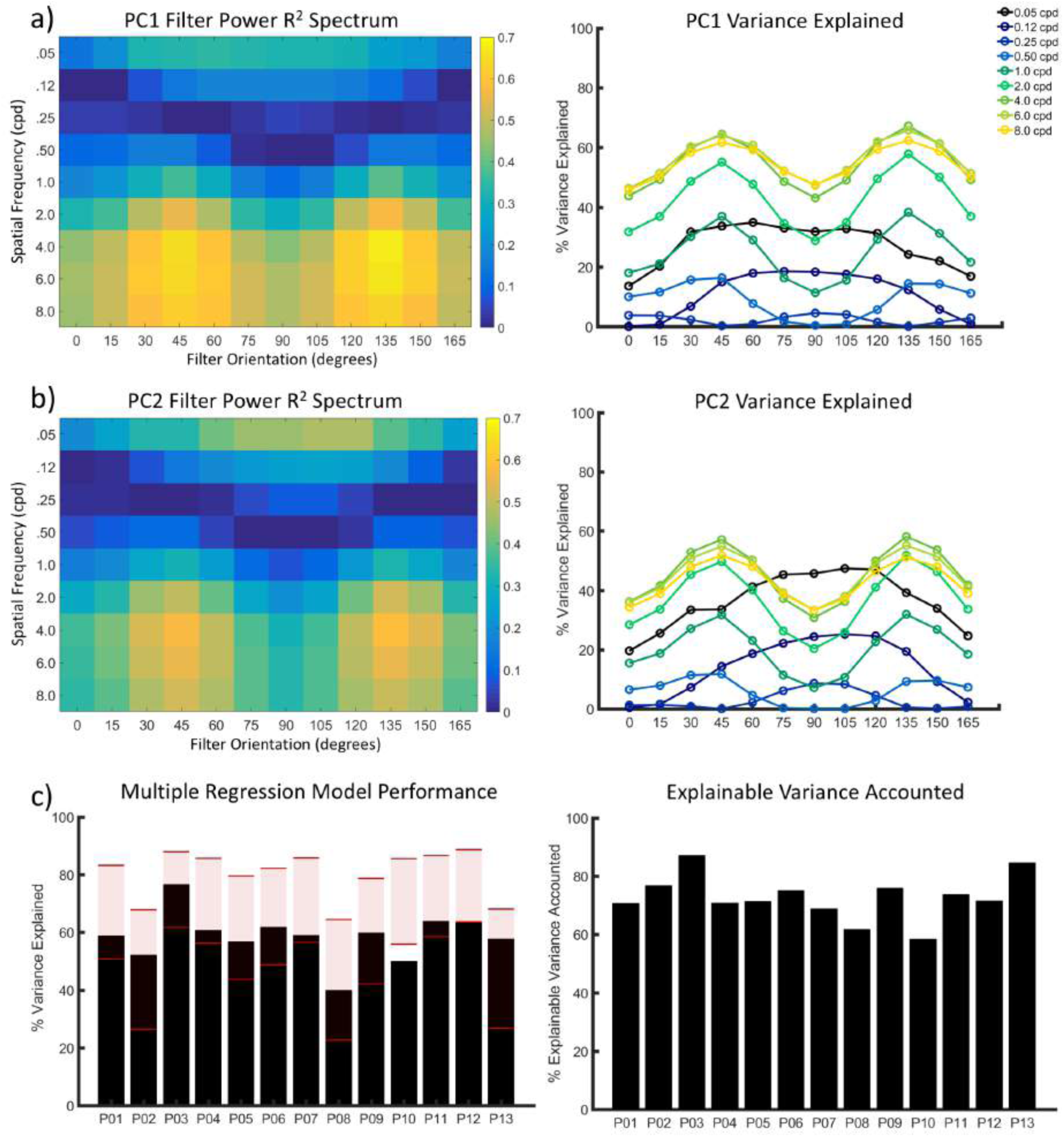
Fourier filter-power model results for Experiment 1. (a) On the left is an R^2^ spectrum showing the regression coefficient for each filter against PC1 from the participant averaged time series data. Rows represent peak SF of the filters in cpd, columns show peak orientation (in degrees) of the filters. On the right are the same data shown in standard line plot format. (b) Same as (a), but for PC2. (c) Plotted on the left is the full regression model (18 filter-power predictors; see text for detail) performance according to variance explained (y-axis) for PC1 for each participant (x-axis). The shaded red area represents the lower and upper bounded region for explainable variance (see text for further detail). On the right shows the same results plotted according to explainable variance accounted for by the full regression model (the ratio of variance explained and the upper bound of estimated explainable variance).

The above analyses were repeated for Experiment 2, first using the participant averaged data (**Figure 10a-b**). Here, the estimated upper and lower bounds of explainable variance were calculated across participants as described in **Appendix 3**. Consistent with Experiment 1, 74% of the explainable variance in PC1 coefficients is accounted for by the higher SFs, with 50% of PC 2’s explainable variance again accounted for by the higher SFs. Next, we conducted the same analysis on a participant-by-participant basis (explainable variance bounds could not be estimated for this analysis) and report the results in **Figure 10c**, which, as expected, shows lower overall variance explained (only one trial per image)^3^.

**Figure 10.**
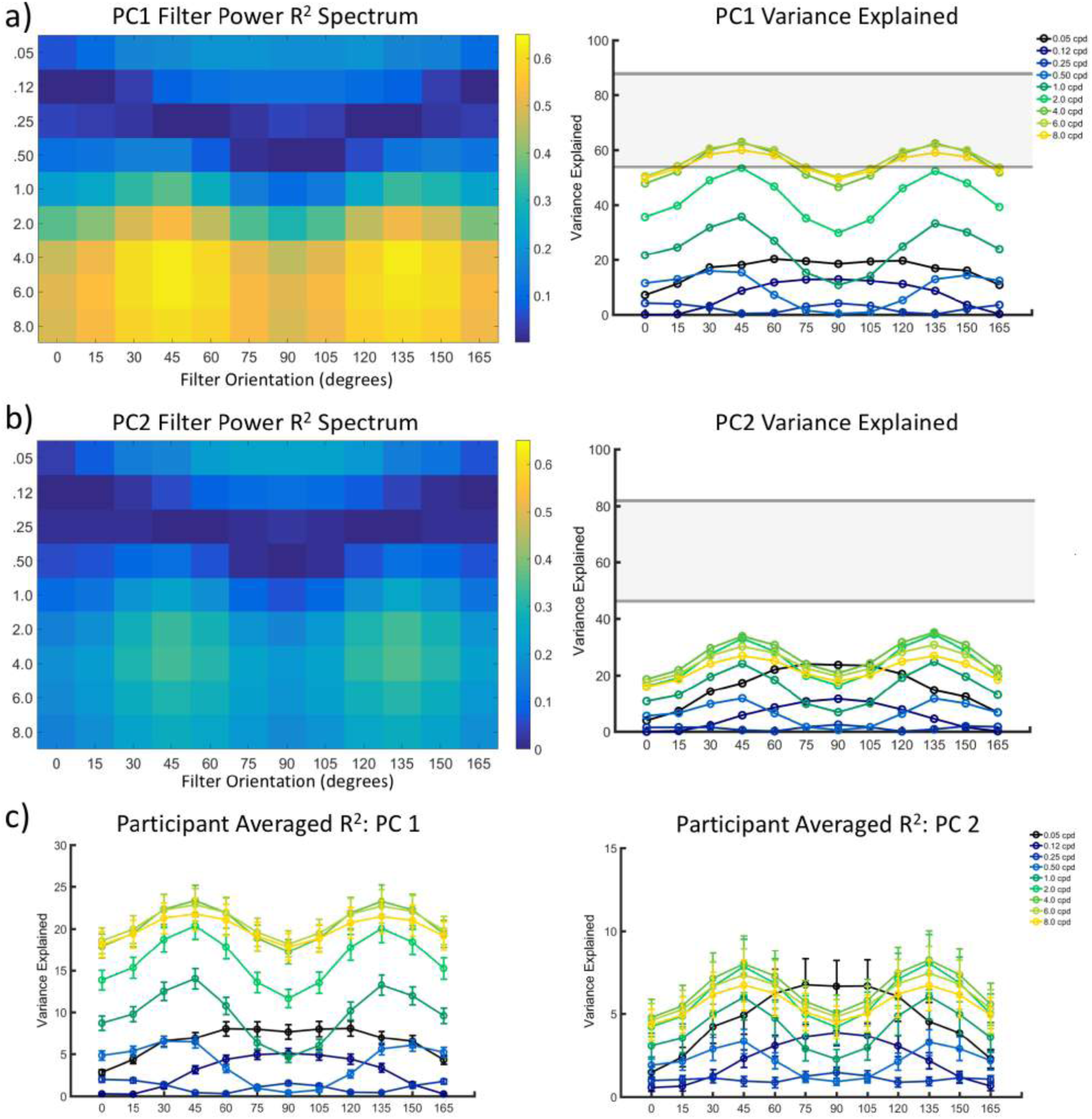
Fourier filter-power model results for Experiment 2. (a) On the left is an R^2^ spectrum showing the regression coefficient for each filter against PC1 from the participant averaged time series data. Rows represent peak SF of the filters in cpd, columns show peak orientation (in degrees) of the filters. On the right are the same data shown in standard line plot format. (b) Same as (a), but for PC2. (c) Plotted on the left is the same as in (a) but was calculated for PC1 of each participant and then averaged (error bars are 95% confidence intervals). The plot on the right was calculated the same way as on the left, but for PC2.

## Experiment 3

The model described above suggests that for both Experiments 1 and 2, the relative positions of images along the two primary PC dimensions of the SSVEP response space can be explained by filter power at SFs ≥ 4 cpd (with the oblique orientations playing a dominant role in that account). However, it is important to note that this model treats each filter as an isolated entity. The difficulty is that the model is looking at only the correlations between SSVEP responses and isolated SF power. Because the stimuli are broadband, correlations with any given frequency band could be due to the simultaneity of that band along with other bands (or some other broadband feature that is correlated with a given band’s power). The current experiment therefore set out to test for a causal explanation for two major observations made from the modeling results. In particular, does the amount of filter power in higher SFs (relative to lower SFs) drive the magnitude of SSVEP SNRs? And, is SSVEP SNR driven more by higher SF oblique orientations than cardinal orientations?

The current experiment consisted of two primary conditions. The first involved measuring SSVEP SNRs while participants viewed bandpass filtered natural images (i.e., narrowband in SF and orientation) that had variable amounts of filter power at each frequency and orientation. If SNRs are being driven by filter power in any given band of SFs and orientations, then we would expect to see SNR increase with increasing filter power within particular bands (e.g., high SF obliques). The second condition was designed to measure the same SNR trends mentioned above, but with compound stimuli. Specifically, all orientations at one SF band would be held constant in terms of Fourier power, while the Fourier power of cardinal or oblique orientations at another SF band are varied. The idea behind this particular stimulus manipulation is to test for an SNR modulation at a band of SFs and orientations due to systematic increases in Fourier power, while simultaneously ‘activating’ neural populations tuned to other SFs and orientations, thereby providing a causal account that is closely related to activity that would likely be observed with broadband stimuli.

### Apparatus

Identical to Experiments 1 and 2, except that the experimental monitor’s gamma was set so that the filtered image pixel values were linear on the display.

### Participants

A total of 16 participants were recruited for this experiment. Of those, 1 failed to produce SNRs at any electrode that exceeded chance SNR (as explained in Experiment 1). The age of the remaining 15 participants (5 female, 13 right-handed) ranged from 18 to 31 (median age = 19). All participants had normal (or corrected to normal) vision as determined by standard ETCRS acuity charts. All participants gave Institutional Review Board-approved written informed consent before participating and were compensated for their time.

### Stimuli & Filtering Procedure

Five natural scene images were pseudo-randomly selected from the stimulus set of Experiment 2. The stimuli were randomly selected from those at the higher end of PC 1 (to optimize SNR) and evaluated to ensure that there was an approximately equivalent amount of Fourier power across all orientations (see [20] for details). The stimuli were then submitted to two linear filtering routines corresponding to the two main conditions of the current experiment. All filtering took place using the log-Gabor functions defined in **Appendix 2**. However, here we reduced edge effects by using the standard symmetrizing technique (e.g., all stimuli were copied and flipped left-to-right, with the result copied and flipped top-to-bottom, thereby doubling the dimensions of the original image) prior to filtering. Target SFs were doubled to account for the increase in dimensions. The original 512 × 512 image was then cropped from the symmetrized filtered image.

For the bandpass filtering condition, all stimuli were filtered to target one of two peak SFs (namely, 0.125 cpd and 4 cpd), with the SF bandwidth of the filter set to 1 octave (full width at half height), and one of four different orientations (vertical, 45° oblique, horizontal, and 135° oblique) with an orientation bandwidth of 16° (full width at half height). Once filtered, 5 copies were made by scaling the power from 12.5 to 14.5 (log units) in steps of 0.5. Thus, each image yielded 40 stimulus images (2 SFs by 4 orientations × 5 levels of power).

For the compound filtering condition, two different sets of filtered images were created. One set contained image content from all orientated filters at 0.125 cpd (1 octave SF bandwidth, 16° orientation bandwidth) combined with either the cardinal orientations (0° *and* 90°) or the oblique orientations (45° *and* 135°) at 4cpd (1 octave SF bandwidth, 16° orientation bandwidth). The power of the 4 cpd filters was scaled as described above and held constant at 13.5 log units for the 0.125 cpd filters. The other set was the opposite, with all four orientations at 4cpd held at a constant power with variable power at 0.125 cpd for the cardinal or oblique orientations. Thus, each image yielded 20 stimulus images (5 overall levels of power × 2 variable orientations at 0.125 × 2 variable power orientations at 4 cpd).

In total, each stimulus image had 60 different filtered versions, for a total of 300 different stimuli.

### Procedure

The experiment consisted of a single recording session lasting ∼48 min. All 300 stimuli were presented once and were randomly interleaved. The trial sequence and temporal contrast modulation were identical to Experiments 1 and 2. Participants engaged in a distractor task that was identical to Experiment 1, except here, there was a luminance change on 60 trials (randomly distributed across the session) and was applied to all filtered stimuli from one of the five images. Participants reported (via gamepad response) when a luminance change occurred and were given the opportunity to rest every 50 trials. The trials that contained a luminance change were not included in the analysis, resulting in a total of 240 stimuli used in analysis.

The EEG recording details, data processing pipeline, and electrode selection routine were identical to Experiments 1 and 2. Artifact trials (no more than 5% across all participants) were found to have no influence on the fundamental frequency and were therefore retained for all subsequent analyses. Thus, each participant produced an electrode-averaged (3 to 8 electrodes selected per participant) time series matrix that was 5000 (time points) × 240 (stimuli). That matrix was then split into the 60 different data blocks described above and averaged across the four exemplar images at each level of Fourier power, yielding a 5000 × 60 time series data matrix for each participant.

### Results

Signal to noise ratios were calculated for the 5Hz fundamental (as described in the previous experiments) for each of the 60 averaged time series waveforms (60 data points for each participant), then averaged across participants and plotted in **Figure 11**. Beginning with the bandpass conditions, Fourier power has virtually no influence on SNR for 0.125 cpd stimuli (**Figure 11a**). This was verified with four different repeated measures ANOVAs, all of which failed to show a main effect of power or significant linear trend (all p’s > 0.05). However, as Fourier power increases for stimuli filtered at 4 cpd (**Figure 11b**), all orientations yield SNRs that increase linearly (significant linear trend; all p’s < 0.022; all partial η^2^s > 0.32), and a significant main effect of power for all orientations (all p’s < 0.031; all partial η^2^s > 0.185). Follow-up paired t-tests revealed no significant differences between the orientation SNRs at higher levels of power. The SNR trends for the compound stimuli show a similar pattern. That is, as power in the 0.125 cpd band for the cardinal or oblique orientations increases against a 4 cpd fixed power pedestal, SNR is virtually unchanged (**Figure 11c**). This observation was confirmed with two different repeated measures ANOVAs, all of which failed to show a main effect of power and no significant linear trend (all p’s > 0.05). This is in direct contrast to the complementary compound condition where SNR increases with increasing power of the 4 cpd band against a 0.125 cpd fixed power pedestal for both orientation sets (**Figure 11d**). Again, this was verified with two different repeated measures ANOVAs that showed a main effect of power (both p’s < 0.03; both partial η^2^s > 0.193), and a significant linear trend (both p’s < 0.023; both partial η^2^s > 0.32). Follow-up paired t-tests failed to differentiate between cardinal and oblique orientations at the higher levels of power.

**Figure 11.**
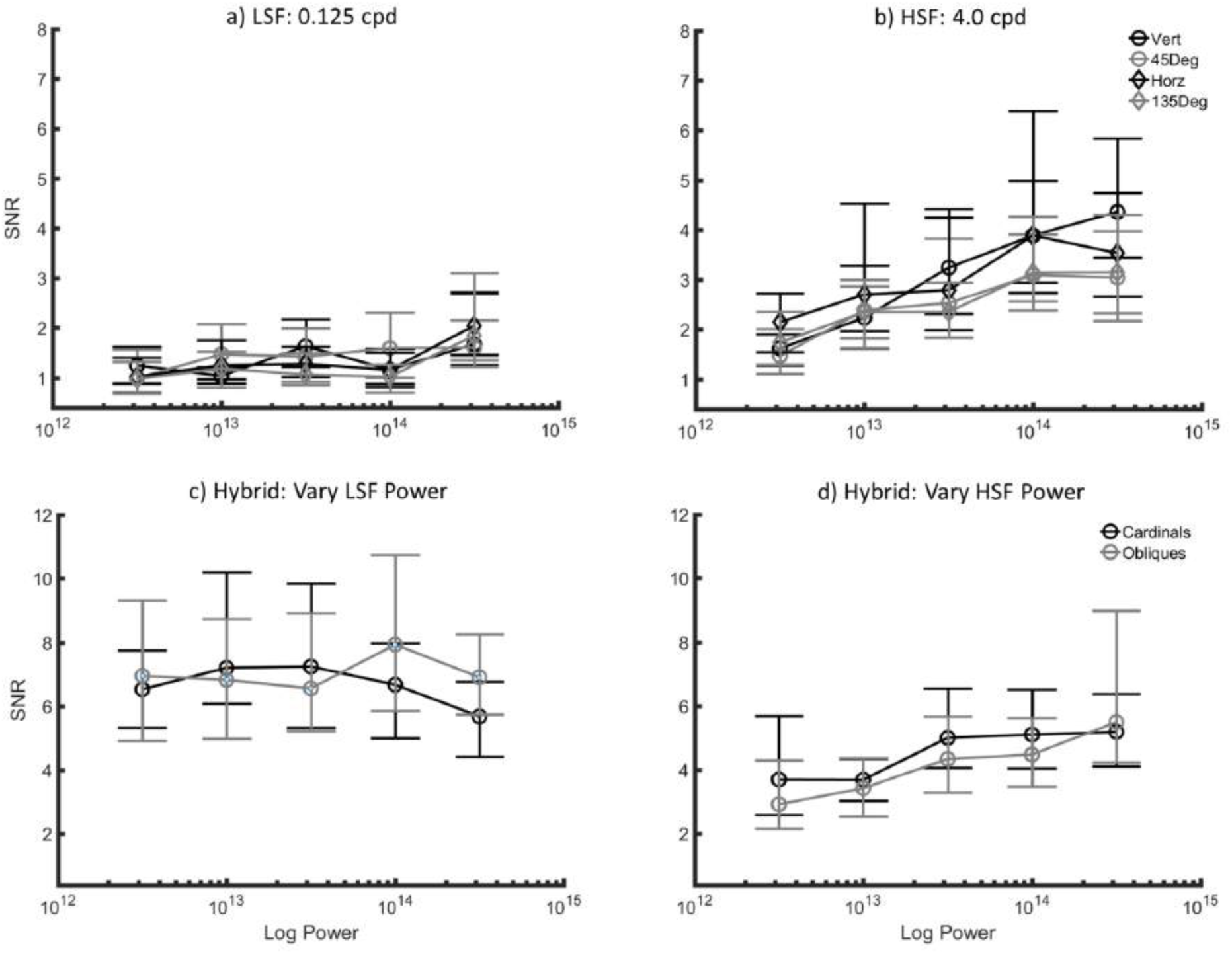
Results from Experiment 3. All plots show SNR for the 5Hz fundamental on the y-axis. The x-axis shows stimulus power for the peak SF and orientation that was modulated for each condition. Refer to the text for further detail.

Together, the results of Experiment 3 support a causal role of power at the higher SFs (≥ 4 cpd) in driving SSVEP signal strength, thereby defining the organization of the SSVEP state-space as measured here. However, we did not observe a prominent role for the oblique orientations, suggesting that the predictive bias observed in the model results of Experiments 1 and 2 most likely resulted from coincidental variation in power at other orientations.

## General Discussion

Considered together, the results of the current study provide a description of the information carried by the SSVEP signal in response to natural scenes, as well as provide a quantitative account of the state-space geometry of the brain responses to those scenes. We believe that this approach has significant advantages for understanding the nature of the SSVEP signal and for understanding what the SSVEP signal can tell us about the processing and organization of natural scenes based on early visual signals.

First, we note that our experiments demonstrate that in response to natural scenes, the SSVEP signal is low dimensional. Although, the Fourier components of the de-noised signal imply that the signal is eight dimensional or fewer, PCA on this signal demonstrates that it is significantly lower. Specifically, the first two PCs of the SSVEP response space captured almost all of the signal’s variance (> 94% across both experiments). The low dimensionality of this response space has both methodological advantages and disadvantages. One important disadvantage is that the low number of dimensions implies that the SSVEP signal cannot capture the vast statistical diversity of natural scenes. On the other hand, with a low-dimensional signal, we can easily visualize the relationships coded by the neural population response and visualize the dependencies of these dimensions. Although most of the variance in the signal is captured by the first PC, the second PC does account for significant variability (∼4% across both experiments), and is well explained by the phase of the SSVEP waveforms. Furthermore, as shown in **Figures 2** and **6**, the two eigenvectors of the response are not independent of each other, which may reflect a dependence of SSVEP phase (i.e., entrainment lag or advance) on image contrast within different bands of spatial frequency.

In the Information Analysis section, we provided an estimate of the information carried by the SSVEP signal regarding our set of natural scenes. As we have noted, our estimates are restricted to the parameters of experiment. Under the conditions of Experiment 1, where all the images are natural scenes with normalized contrast, and using test/retest reliability, we find that the SSVEP provides ∼2.1 bits of information about the image. We believe that although this this number may seem relatively low (i.e, it allows us to categorize these scenes in just over 4 categories), this is a useful approximation. However, that number is based on a set of assumptions regarding the inherent noise in test/retest reliability and how representative our images are with respect to the population of natural scene images. We are currently investigating some of these assumptions, but we believe this approach can provide important insight into both SSVEP and VEP signals.

A more interesting approach is to consider the nature of the information that appears in the population response. In the modeling section, we used a biologically plausible neural model of activity where the images were compared through the sum of squared filter responses. This is basically a complex cell model where the SSVEP signal represents coarse information regarding orientation and SF. Subsequent modeling of the two dominant SSVEP state-space dimensions revealed that filter-power could be used to explain the relative positions of image signals along each PC dimension of the SSVEP neural response space. Interestingly, filter-power at SFs ≥ 4 cpd played a dominant role in defining the organization of images within the neural state-space. Further, collapsing the filter bank to cardinal and oblique orientations across nine SFs (just 17% of all filters initially employed) could explain ∼73% of PC1 and 60% of PC2. We were surprised at the relative power of this simple model. Using machine learning techniques, we may be able to improve on this explanatory power. However, the filter-power model provides a more straightforward interpretation of the organization of images based on SSVEP signals. Indeed, filter-power models have been successful in explaining relatively large amounts of response variance in other studies focused on natural image processing using different neural recording techniques [e.g., 16, 43-45].

Another interesting facet of the SSVEP state-space concerns the role that higher SFs play in defining the relative positions of image responses in that space. We see a similar response dependency in the visual evoked potential (VEP) literature. Specifically, the earliest VEP measured for stimuli presented to the fovea (the fC1, [42]) shows a highpass tuning response for sinusoidal grating stimuli beginning around 4 cpd and increasing in magnitude with increasing SF [42, 46–48]. The latency of that component also increases with increasing SF from ∼75 msec out to ∼90 msec but has been observed as late as 120 msec [48]. Further, when systematically extending sinusoidal gratings (or narrowband filtered natural images) to broadband natural images, the magnitude of that component as well as a later negativity peaking around 150 msec is driven by RMS contrast at the higher SFs [14–15]. While the current study was not designed to map the mechanisms that were entrained by our SSVEP paradigm to those responsible in signaling components in VEP paradigms, the connection between the current results and those from VEP studies using natural scenes is intriguing. Interestingly, data that we collected for another study (manuscript in preparation) suggest that the VEP response at 100-200 msec post stimulus onset is highly correlated (maximum R^2^ = 0.78) with PC1 in the SSVEP response. This is the VEP time range that typically shows selectivity to high SF contrast in natural images (or sinusoidal gratings) [14–15] and suggests that the mechanisms responsible for those VEPs are those largely entrained by our SSVEP paradigm.

The reliance on high SFs observed here (as well as in the VEP literature) has several interesting implications for cortical whitening. It has been argued that in response to the 1/f structure of natural scenes, the visual system increases the relative gain of higher versus lower spatial frequencies [49–50]. Such an approach would explain why white noise perceptually appears to be dominated by high SFs. However, while the apparent role of high SFs in shaping the neural state-space is interesting, it is important to temper those claims given the coarse nature of EEG recordings. That is, EEG is only able to measure the largest neural responses arising from piecemeal cortical signals (e.g., those best aligned to yield dipolar summation), and will not pick up signals from underlying populations that are too weak or happen to cancel out due to the relative orientation of the pyramidal cell generators. For example, the more peripheral signals may cancel due to their origins in the upper and lower banks of the calcarine, thereby leaving only activity arising from the fovea, which would be dominated by high SF responses [51–54]. Thus, the dominance of high SFs may be due to an important transformation of broadband input but could also simply result from being a signal that largely arises from foveal generators. We are currently running experiments in order to advance one hypothesis over the other.

In sum, we have described a state-space approach for characterizing the SSVEP response to natural scenes. We have found that the signal is low dimensional but that the signal contains significant information regarding the image content. We believe this approach provides important insights into the information carried by this signal. We are currently using this approach to investigate the geometry and dimensionality of the VEP signal. Although we find this non-entrained signal to be more complex, the signal is still relatively low dimensional (98% of the variance captured in the first 4 dimensions). We believe that describing the response in terms of the location in this high dimensional space is much more informative than current approaches that focus on the positions and amplitudes of particular features.

The early geometric representation of natural images serves as an important step to elucidate “higher” level state-space representations. One possible candidate model for bridging the current findings to higher knowledge spaces may be the spatial envelope model [55]. In fact, the feature vectors that contribute to the linear discriminant filters (i.e., filters built from LDA) in that model consist of Gabor wavelet responses (similar to those used here) to natural scenes. While LDA may not be a viable means for modeling the transformation of an early filter space to a high categorical space, the initial geometric representation of the input to that model certainly seems to be plausible given the model performance observed in the current study.

### Conclusion

In this study, we have described the information carried by the SSVEP signal in response to natural scenes, as well as the state-space geometry of the responses to those scenes. Further, we show that a log-Gabor filter model can account for a high proportion of the explainable variance along each axis of the SSVEP state-space (∼73% for PC1 and ∼60% for PC2). This suggests that the majority of the information in the entrained neuronal signal can be explained by lower-level statistical features of the stimuli (high SF filter responses in particular). It may well be that higher percentages of the variance could be explained by models that also include higher level features that are not fully explained by the lower level features (emotional impact, task relevance, etc.). However, the success of the feature-based model implies that low-level features play the primary role in the SSVEP entrainment. An understanding of how the electric dipoles are produced by the millions of active neurons, is beyond the scope of this paper. However, this paper provides clear evidence that there is significant information about natural scenes contained in this non-invasive signal.

We believe that this approach can also be applied to VEP data and we are currently exploring that option. From our geometric point of view, each response (VEP or SSVEP) is a point in the relatively low dimensional space of possible responses. By understanding the geometric distribution of these points, we believe we can gain important insights into the nature of the transformation of the visual signal. Further work is needed to understand how particular features of the stimulus influence the position of each image in this space. However, this approach allows us to characterize the full response to individual stimuli in a relatively low dimensional space and does not simply focus on particular features of that space. Finally, the state-space approach does not depend on the particular recording technique that is used (e.g., EEG, MEG, fMRI, fNIRS), and lends itself to any type of encoding model. In fact, recent fMRI work has used PCA to build state-space like response spaces from lexical encoders [e.g., 58]. All recording techniques will result in some type of response for each stimulus. Projecting those responses into a state-space framework as we have done here will put them in a common metric space, enabling direct comparision of response spaces built from common sets of stimuli. Such comparisons may enable insight into how different neural circuits are contributing to the signals captured by macro-scale measures such as those mentioned above.

## Acknowledgements

This work was funded by a National Science Foundation grant (1736274) to BCH and MRG, James S McDonnell Foundation grant (220020439) to BCH, and a Colgate Summer Undergraduate Fellowship to CO.

## Appendix

### 1.0 RMS contrast manipulations

Root mean square contrast is defined as the standard deviation of all pixel luminance values divided by the mean of all pixel luminance values. Here, images are treated as arrays, I(y), with each array set to have the same RMS contrast and zero mean using the following operations.

The pixels values of each image array are first normalized to fall between [-.5 .5] with zero mean as follows,

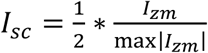

with I_zm_ defined as:

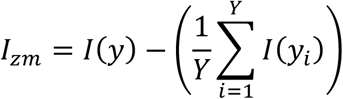

Here, Y represents the total number of pixels in each array. Next, we calculated RMS for I_sc_ as follows:

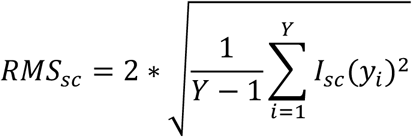

We then calculated an RMS scaling factor, S_rms_ = (2*RMS_t_)/RMS_sc_, with RMS_t_ set to a reasonable target RMS value. By reasonable, we mean a value that did not result in significant (> 5%) clipping of the resulting pixel values. That value was 0.20 for the images used in the current study. Finally, each image array was scaled to have an RMS equal to RMS_t_ and reassign to I(y) as follows: I(y) = 127*(I_sc_*S_rms_). Note that scaling by 127 puts the scaled pixel values of I(y) back in the original range of I_zm_. Image arrays were then converted back to matrix form.

### 2.0 Measuring log-Gabor filter power

All image filtering was conducted in the Fourier domain using the images in matrix form. The images were first made to possess a zero mean and an RMS contrast of 0.20 (see Appendix 1). To minimize edge effects in the Fourier domain due to the non-periodic nature of natural images, the images were multiplied with a 2D circular Hann window prior to the Fast Fourier transform. To construct the window, a 2D radial matrix, M_R_, was built within which the values increased from the center out to 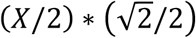 and modulated according to:

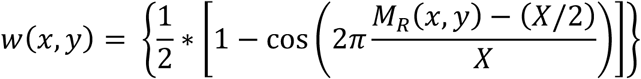

with X being the maximum dimension of the image to be weighted by w(x,y). Each windowed image was submitted to the 2D fast Fourier transform to obtain H(u,v) as follows:

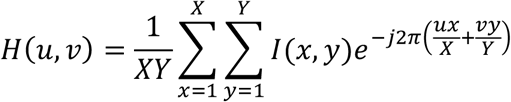

where I(x,y) represents a given image, with X and Y representing the dimensions of the image. Next, the amplitude spectrum was calculated according to:

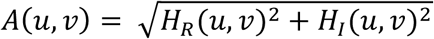

with H_R_(u,v) and H_I_(u,v) representing the real and imaginary parts of H(u,v), respectively. For filtering convenience, the amplitude spectrum, A(u,v) was shifted to polar coordinates and in this form will be denoted as A(*f*,θ), with *f* serving as the index along the radial (i.e., spatial frequency) dimension, and θ as the index along the theta (i.e., orientation) dimension.

A 2D log-Gabor filter [46] in the Fourier domain consists of a log-Gaussian function along the *f* axis and a Gaussian function along the θ axis, and can be obtained by multiplying a 2D log-Gaussian filter (i.e., a log ‘doughnut’ filter) with a 2D Gaussian ‘wedge’ filter. The construction of the 2D log-Gaussian filter, L_gaus_(*f*, θ), took place in same polar coordinate frame as A(*f*,θ). Thus, for each θ axis, L_gaus_(*f*) was modulated as follows.

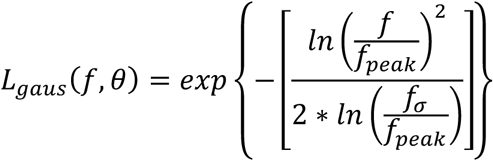

Where *f* increases with spatial frequency (radial distance), *f*_*peak*_ represents the peak of the function, and *f*_*σ*_ representing the SF bandwidth of the filter. Next, a 2D Gaussian function (modulated across θ in radians) about a central orientation was generated as follows.

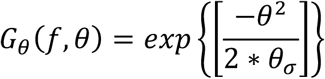

The log-Gabor filter, LG(*f*, θ), was then constructed by multiplying G_θ_(*f*, θ) by L_gaus_(*f*, θ). To obtain filter power, the amplitude spectrum in polar coordinates, A(*f*, θ), was first squared to obtain the power spectrum, P(*f*, θ). Power for each filter then measured as follows.

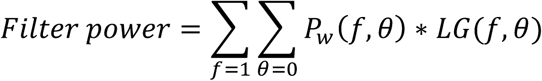

Where P_w_(*f*,θ) is the power spectrum multiplied by a hard-edged circular window containing ones everywhere within a diameter equal to the image dimension and zeroes everywhere else. Applying that window ensured equal SF sampling across all orientations.

### 3.0 Explainable Variance Bounds

The upper and lower bounds of the model regression coefficients, R^2^, were calculated according to the following expression (based on [56–57]).

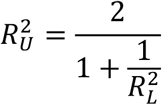

Here, the upper bound, R_U_^2^, is the estimated R^2^ that would be expected between PCs taken from the actual neural signal (i.e., as averaged over an infinite number of samples, thereby containing zero noise) and the observed PCs (i.e., PCs extracted from a finite number of averaged trials). This value therefore represents an approximation to the highest R^2^ that would be expected given the noise inherent in the data. The lower bound, R_L_^2^, is the averaged R^2^ taken between pairs of PCs extracted from the average of pairs of trial repetitions (Experiment 1) or the average across half of the participants compared to the other half (Experiment 2). Specifically, for Experiment 1 an R_L_^2^ was measured for each participant by calculating R^2^ between a given PC based on the average of two trial repetitions and the same PC from the average of the other two repetitions. This process was repeated for all possible unique pairs of repetitions and then averaged. For Experiment 2, R_L_^2^ was measured by calculating R^2^ between a given PC from the averaged data of half of the participants and the same PC form the average of the other half of the participants. This process was repeated 100 times (each time randomly assigning different participants to the first and second half averages) and the average of those R^2^s was taken as R_L_^2^.

## Supplementary Material

**Figure S1.**
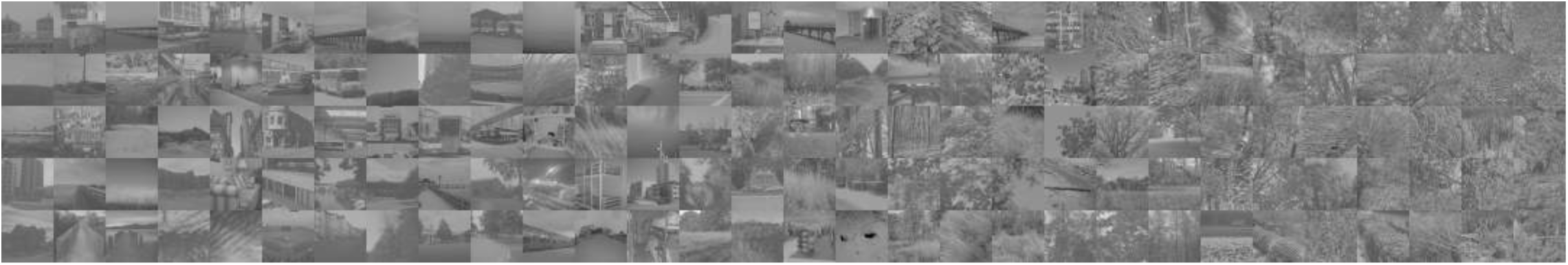
Experiment 1. Stimuli binned along PC 2 (x-axis of the image array). To facilitate a visual presentation, the images were sorted according to the participant-averaged PC 2 coefficients and binned into 5 sets with 30 images per bin (thus there is no inherent meaning to the y-axis of the image array).

**Figure S2.**
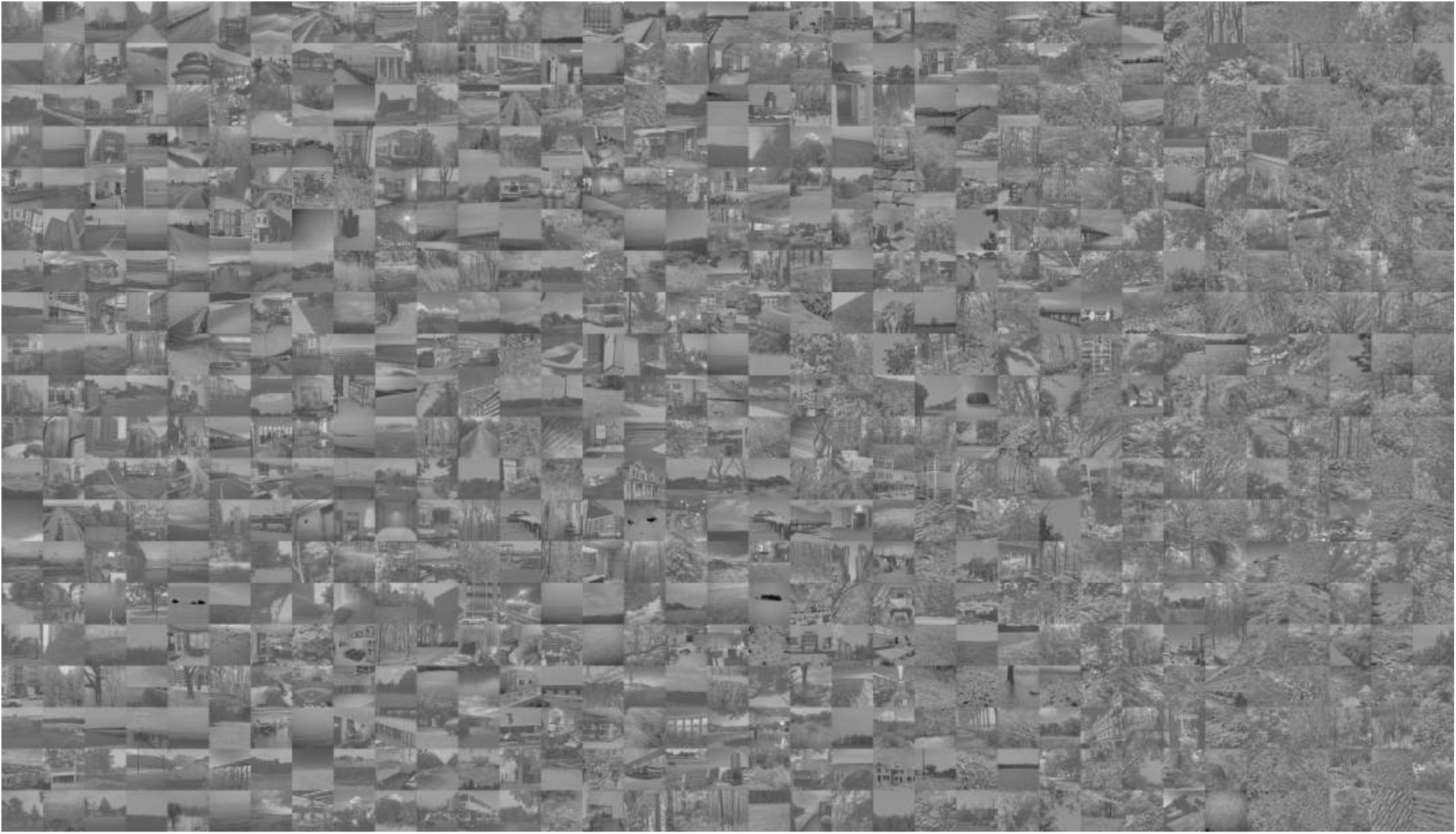
Experiment 2. Stimuli binned along PC 2. To facilitate a visual presentation, the images were sorted according to the participant-averaged PC 2 coefficients (low to high) and binned into 7 sets with 100 images per bin (as with **Figure S1**, row membership is arbitrary).

## Footnotes

1 We chose 5 Hz because it enabled a smooth sinusoidal temporal profile (yet was high enough to allow several cycles of oscillation) and was sufficiently below the 10 Hz alpha bias known to dominate occipital electrodes. Our measures of SSVEP state-space are therefore specific to that particular stimulus modulation and might not generalize well to higher stimulus modulation frequencies that may recruit neural populations with different temporal sensitivities.

2 We tested grid sizes between 2×2 to 500×500, and found stable results for grid sizes larger than 50×50.

3 We tested the explanatory power of the filter-power encoding model relative to other image encoding models based on well-established image statistics (e.g., amplitude spectrum slope, structural complexity, orientation bias, whitened skewness, whitened kurtosis, and the slope of the phase-only second spectrum). To cut down on the number of predictors associated with the filter-power model, all responses were averaged over orientation (i.e, 9 predictors in total for each image). Multiple regression was run on each of the PC coefficients from Experiment 1 with the outputs of the reduced filter-power model and the the six image statistic models used as predictors). The high spatial frequencies (>= 4 cpd) of the filter-power model explained more *unique variance* for each of the PC coefficients (25% and 17% represpectively) than any of the image statistics encoder models (highest predicted unique variance across all six image encoder models = PC1: 8%; PC2: 3%).

## References

1. Baddeley, R. Abbott, L.F., Booth, F.S., Freeman, T., Wakeman, E.A., & Rolls, E.T. (1997). Reponses of neurons in primary and inferior temporal visual cortices to natural scenes. Proceedings of the Royal Society London B, 264, 1775–1783.

2. Dan, Y., Atick, J.J., & Reid, R.C. (1996). Efficient coding of natural scenes in the lateral geniculate nucleus: Experimental test of computational theory. Journal of Neuroscience, 16, 3351–3362.

3. David, S.V., Vinje, W.E., & Gallant, J.L. (2004). Natural stimulus statistics alter the receptive field structure of V1 Neurons. Journal of Neuroscience, 24, 6991–7006.

4. David, S.V. & Gallant, J.L. (2005). Predicting neuronal responses during natural vision. Network: Computation in Neural Systems, 16, 236–260.

5. Felsen, G., Touryan, J., Han, F., & Dan, Y. (2005). Cortical sensitivity to visual features in natural scenes. PLoS Biology, 3, 1819–1828.

6. Freeman, J., Ziemba, C.M., Heeger, D.J., Simoncelli, E.P., & Movshon, J.A. (2013). A functional and perceptual signature of the second visual area in primates. Nature Neuroscience, 16, 974–981.

7. Mante, V., Frazor, R.A., Bonin, V., Geisler, W.S., & Carandini, M. (2005). Independence of luminance and contrast in natural scenes and in the early visual system. Nature Neuroscience, 12, 1690–1697.

8. Tolhurst, D.J., Smyth, D., & Thompson, I.D. (2009). The sparseness of neuronal responses in ferret primary visual cortex. Journal of Neuroscience, 29, 2355–2370.

9. Weliky, M., Fiser, J., Hunt, R.H., & Wagner, D.N. (2003). Coding of natural scenes in primary visual cortex. Neuron, 37, 703–718.

10. Ayzenshtat, I., GIlad, A., Zurawel, G., & Slovin, H. (2012). Population response to natural images in the primary visual cortex encodes local stimulus attributes and perceptual processing. Journal of Neuroscience, 32, 13971–13986.

11. Kayser, C., Salazae, R.F., & König, R. (2003). Responses to natural scenes in cat V1. Journal of Neurophysiology, 90, 1910–1920.

12. Tang, S., Zhang, Y., Li, Z., Li, M., Liu, F., Jiang, H., & Lee, T S. (2018). Large-scale two-photon imaging revealed super-sparse population codes in the V1 superficial layer of awake monkeys. eLife, 7, e33370.

13. Groen, I.I.A., Ghebreab, S., Prins, H., Lamme, V.A.F., & Scholte, H.S. (2013). From image statistics to scene gist: Evoked neural activity reveals transition from low-level natural image structure to scene category. Journal of Neuroscience, 33, 18814–18824.

14. Hansen, B.C., Jacques, T., Johnson, A.P., & Ellemberg, D. (2011). From spatial frequency contrast to edge preponderance: The differential modulation of early visual evoked potentials by natural scene stimuli. Visual Neuroscience, 1–12.

15. Hansen, B.C., Johnson, A.P., & Ellemberg, D. (2012). Different spatial frequency bands selectively signal for natural image statistics in the early visual system. Journal of Neurophysiology, 108, 2160–2172.

16. Kay, K.N., Naselaris, T., Prenger, R.J., & Gallant, J.L. (2008). Identifying natural images from human brain activity. Nature, 452, 352–356.

17. Nguyen, T., Kuntzelman, K., & Miskovic, V. (2017). Entrainment of visual steady-state responses is modulated by global spatial statistics. Journal of Neurophysiology, 118, 344–352.

18. Nishimoto, S., Vu, A.T., Naselaris, T., Benjamini, Y., Yu, B., & Gallant, J.L. (2011). Reconstructing visual experiences from brain activity evoked by natural movies. Current Biology, 21, 1641–1646.

19. Bex, P. J., Solomon, S. G., & Dakin, S. C. (2009). Contrast sensitivity in natural scenes depends on edge as well as spatial frequency structure. Journal of Vision, 9(10), 1 1–19.

20. Hansen, B.C. & Essock, E.A. (2004). A horizontal bias in human visual processing of orientation and its correspondence to the structural components of natural scenes. Journal of Vision, 4, 1044–1060.

21. Long, F., Yamng, Z., & Purves, D. (2006). Spectral statistics in natural scenes predict hue, saturation, and brightness. Proceedings of the National Academy of Sciences, 103, 6013–6018.

22. Tadmor, Y., & Tolhurst, D. J. (1994). Discrimination of changes in the second-order statistics of natural and synthetic images. Vision Research, 34(4), 541–554.

23. Webster, M. A., & Miyahara, E. (1997). Contrast adaptation and the spatial structure of natural images. Journal of the Optical Society of America A, 14(9), 2355–2366.

24. Yang, Z. & Purves, D. (2003). A statistical explanation of visual space. Nature Neuroscience, 6, 632–640.

25. Norcia, A.M., Appelbaum, L.G., Ales, J.M., Cottereau, B.R., & Rossion, B. (2015). The steady-state visual evoked potential in vision research: A review. Journal of Vision,15, 1–46.

26. Regan, D. (1989). Human brain electrophysiology: Evoked potential and evoked magnetic fields in science and medicine. Amsterdam, the Netherlands: Elsevier.

27. Regan, D. (1966). Some characteristics of average steady-state and transient responses evoked by modulated light. Electroencephalography and Clinical Neurophysiology, 20, 238–248.

28. Regan, D. & Regan, M.P. (1988). A frequency domain technique for characterizing nonlinearities in biological systems. Journal of Theoretical Biology, 133, 293–317.

29. Field, D.J. (1994). What is the goal of sensory coding? Neural Computation, 6, 559–601.

30. Golden, J.R., Vilankar, K.P., Wu, M.C.k., & Field, D.J. (2016). Conjectures regarding the nonlinear geometry of visual neurons. Vision Research, 120, 74–92.

31. Zetzsche, C. & Nuding, U. (2005). Nonlinear and higher-order approaches to the encoding of natural scenes. Network, 16, 191–221.

32. Greene, M.R. & Hansen, B.C. (2018). Shared spatiotemporal category representations in biological and artificial deep neural networks. PLoS Computational Biology, 14, e1006327.

33. Xiao, J., Ehinger, K.A., Hays, J., Torralba, A., & Oliva, A. (2014). SUN database: Exploring a large collection of scene categories. International Journal of Computer Vision, DOI 10.1007/s11263-014-0748-y

34. Hansen, B.C. & Hess, R.F. (2006). Discrimination of amplitude spectrum slope in the fovea and parafovea and the local amplitude distributions of natural scene imagery. Journal of Vision, 6, 696–711.

35. Berens, P. (2009). Circstat: A matlab toolbox for circular statistics. Journal of Statistical Software,31, 1–21.

36. Torralba, A. & Oliva, A. (2003). Statistics of natural image categories. Network: Computation in Neural Systems, 14, 391–412.

37. Chandler, D.M. & Field, D.J. (2007). Estimates of the information content and dimensionality of natural scenes from proximity distributions. Journal of the Optical Society of America A, 24, 922–941.

38. Carandini, M., Demb, J.B., Mante, V., Tolhurst, D.J., Dan, Y., Olshausen, B.A., Gallant, J.L., & Rust, N.C. (2005). Do we know what the early visual system does? Journal of Neuroscience, 25, 10577–10597.

39. De Valois, R.L., Albrecht, D.G., & Thorell, L.G. (1982). Spatial frequency selectivity of cells in macaque visual cortex. Vision Research, 22, 545–559.

40. De Valois, R.L., Yund, E.W., & Hepler, N. (1982). The orientation and direction selectivity of cells in macaque visual cortex. Vision Research, 22, 531–544.

41. Kriegeskorte, N., Mur, M., & Bandettini, P. (2008). Representational similarity analysis – Connecting the branches of systems neuroscience. Frontiers in Systems Neuroscience, 2, 1–28.

42. Hansen, B.C., Haun, A.M., Johnson, A.P., & Ellemberg, D. (2016). On the differentiation of foveal and peripheral early visual evoked potentials. Brain Topography, 29, 506–514.

43. Nishimoto, S., Vu, A.T., Naselaris, T., Benjamini, Y., Yu, B, & Gallant, J.L. (2011). Reconstructing visual experiences from brain activity evoked by natural scenes. Current Biology, 21, 1641–1646.

44. David, S.V. & Gallant, J.L. (2005). Predicting neuronal responses during natural vision. Network: Computation in Neural Systems, 16, 239–260.

45. Ramkumar, P., Hansen, B.C., Pannasch, S., & Loschky, L.C. (2016). Visual information representation and rapid-scene categorization are simultaneous across cortex: An MEG study. NeuroImage, 134, 295–304.

46. Ellemberg, D., Hammarrenger, B., Lepore, F., Roy, M. S., & Guillemot, J. P. (2001). Contrast dependency of VEPs as a function of spatial frequency: the parvocellular and magnocellular contributions to human VEPs. Spatial Vision, 15, 99–111.

47. Tobimatsu, S., Tomoda, H., & Kato, M. (1995). Magnocellular and parvocellular contributions to visual evoked potentials in humans: Stimulation with chromatic and achromatic gratings and apparent motion. Journal of the Neurological Sciences, 134, 73–82.

48. Vassilev, A., Mihaylova, M., & Bonnet, C. (2002). On the delay in processing high spatial frequency visual information: reaction time and VEP latency study of the effect of local intensity of stimulation. Vision Research, 42, 851–864.

49. Field, D.J. (1987). Relations between the statistics of natural images and the response properties of cortical cells. Journal of the Optical Society of America A, 4, 2379–2394.

50. Brady, N. & Field, D.J. (1995). What’s constant in contrast constancy? The effects of scaling on the perceived contrast of bandpass patterns. Vision Research, 35, 739–756.

51. Henriksson, L., Nurminen, L., Hyvärinen, A., & Vanni, S. (2008). Spatial frequency tuning in human retinotopic visual areas. Journal of Vision, 8, 1–13.

52. Hess, R.F., Li, X, Mansouri, B., Thompson, B., & Hansen, B.C. (2009). Selectivity as well as sensitivity loss characterizes the cortical spatial frequency deficit in amblyopia. Human Brain Mapping, 30, 4054–4069.

53. Sasaki, Y., Hadjikhani, N., Fischl, B., Liu, A.K., Marrett, S., Dale, A.M., & Tootell, R.B.H. (2001). Local and global attention are mapped retinotopically in human occipital cortex. Proceedings of the National Academy of Science, 98, 2077–2082.

54. Singh, K.D., Smith, A.T., & Greenlee, M.W. (2000). Spatiotemporal frequency and direction sensitivities of human visual areas measured using fmri. NeuroImage, 12, 550–564.

55. Oliva, A. & Torralba, A. (2001). Modeling the shape of the scene: A holistic representation of the spatial envelope. International Journal in Computer Science, 42, 145–175.

56. Hsu, A., Borst, A., & Theunissen, F.E. (2004). Quantifying variability in neural responses and its application for the validation of model predictions. Network: Computation in Neural Systems, 15, 91–109.

57. Schoppe, O., Harper, N.S., Willmore, B.D.B., King, A.J., & Schnupp, J.W.H. (2016). Measuring the performance of neural models. Frontiers in Computational Neuroscience, 10:10.

58. Huth, A.G., Nishimoto, S., Vu, A.T., & Gallant, J.L. (2012). A continuous space describes the representation of thousands of object and action categories across the human brain. Neuron, 76, 1210–1224.

